# Small dams drive *Anopheles* abundance during the dry season in a high malaria burden area of Malawi

**DOI:** 10.1101/2023.11.14.567025

**Authors:** Kennedy Zembere, Christopher M Jones, Rhosheen Mthawanji, Clinton Nkolokosa, Richard Kamwezi, Patrick Ken Kalonde, Michelle C Stanton

**Author notes:** These authors contributed equally.

## Abstract

This study explores the influence of small dams on the exposure to malaria vectors during the dry season in Kasungu district, Malawi, an area recently identified as high priority for malaria interventions by the National Malaria Control Programme. Small dam impoundments provide communities with a continuous supply of water for domestic and agricultural activities across sub-Saharan Africa and are considered vital to food security and climate change resilience. However, these permanent water bodies also create ideal breeding sites for mosquitoes in typically arid landscapes. The study focuses on a specific dam impoundment and its vicinity, aiming to assess its spatial and temporal influence on indoor vector densities.

Throughout May to August 2021, CDC light traps were used to measure indoor mosquito densities for two consecutive nights per month in three communities located at increasing distances from the dam (0km, ∼1km, ∼2km). Simultaneously, drone imagery was captured for each community, enabling the identification of additional standing water within approximately 400 meters of selected households. Larval sampling was carried out within the impoundment periphery and in additional water bodies identified in the drone imagery. Generalised linear mixed models (GLMMs) were employed to analyse the indoor *Anopheles* abundance data, estimating the effects of household structure (open/closed eaves), month, temperature, and water proximity on malaria vector exposure.

Throughout 685 trapping nights, a total of 1,256 mosquitoes were captured, with 33% (412) being female *Anopheles*. Among these, 91% were morphologically identified as *An. funestus* s.l., and 5% as *An. gambiae* s.l. Catches progressively declines in each consecutive trapping month as the environment became drier. This decline was much slower in Malangano, the community next to the dam, with abundance being notably higher in June and July. Further, the majority of *An. gambiae* s.l. were caught in May, with none identified in July and August. *Anopheles* larvae were found both in the impoundment and other smaller water bodies such as irrigation wells in each survey month, however the presence of these smaller water bodies did not have a significant impact on adult female mosquito catches in the GLMM. The study concludes that proximity to the dam impoundment was the primary driver of differences between survey communities with the abundance in Chikhombwe (∼1km away) and Chiponde (∼2km away) being 0.35 (95% CI 0.19-0.66) and 0.28 (95% CI 0.16-0.47) lower than Malangano respectively after adjusting for other factors.

These findings underscore the importance of targeted interventions, such as larval source management or housing improvements, near small dams to mitigate malaria transmission risks during the dry season. Further research is needed to develop cost-effective strategies for vector control within and around these impoundments.

## Introduction

The seasonal fluctuation of *Anopheles* mosquito populations is inextricably linked with rainfall. During the rainy season substantial areas of standing water are available for mosquito egg-laying and larval development. Should these habitats persist long enough to exceed the temperature-dependent time it takes for adult emergence, then lagged malaria transmission will inevitably follow in endemic areas. While there is a positive association between rainfall and mosquito density^1,2^, the process is highly non-linear and dependent on local topography, hydrology and rainfall patterns (e.g. distribution and severity of rainfall events)^3^ as well as the ecology of the local dominant species. Once the rains subside, however, mosquito habitat gradually succumbs to evapotranspiration. In certain areas (e.g. the Sahel), this leaves virtually no extant pools for oviposition, causing *Anopheles* populations to enter either a state of dormancy (aestivation)^4^ or undergo wind-assisted migration^5^ to maintain population viability. That said, in many parts of Africa, enough standing water persists into the dry season extending the malaria transmission season, disproportionately increasing the risk of those communities living in proximity to dry season habitat.

Natural river and lakeside habitat may provide suitable dry season habitat for mosquito production, but most remaining water bodies are invariably tied to human practices and particularly those related to agriculture and water security. This creates a dichotomy between socio-economic development and the risk of infectious disease^6^. Permanent and semi-permanent water holding structures such as irrigation canals, water impounded by dams, urban agriculture, and water storage units, all promote *Anopheles* production in the absence of rainfall^7^. Epidemiological and entomological observations surrounding individual dams repeatedly show an increased risk of malaria in those people living in proximity to the dam periphery, and these observations are supported by larger systematic data analyses across the continent. Most of the research on dams and malaria to date has focussed on large dams^8^. These are defined as those constructions with a height of 15 metres resulting in impoundments holding 3 million m^3^ of water. The relative contribution of earthen small (or micro) dams – only a few metres high with impoundments containing 100-500,000 m^3^ of water - on malaria incidence may in fact be greater. Small dam impoundments provide a more suitable habitat for *Anopheles* due to their shallower and more vegetated edges^7^. Furthermore, communities tend to live in closer proximity to these structures with greater accessibility and use the water directly for livestock watering and domestic activities.

Malawi has a predominantly agriculture-based economy which depends on increasingly erratic annual rainfall patterns^9^. Small dams form part of Malawi’s growing efforts to double the area of land under irrigation by 2035^10^. They provide some security in terms of water accessibility and are found across Malawi, however no current database of the location, size and status of these dams exists^11^. The general interplay between irrigation and malaria in Africa is well-understood but the relationship is complex and depends on the agroecosystem, socio-economic factors, local vector behaviour and ecology, and the seasonality of disease^12,13^. Efforts to characterise this relationship in Malawi have been performed in a commercial sugar growing estate and within an irrigated rice valley^14,15^, and research is underway to do the same within a World Bank funded large-scale transformation programme^16^. These systems are, however, much larger and concentrated on a single geographic area. Small dams are different in that their impoundments are much smaller and their impact on mosquito abundance and disease is likely to be far more focal.

The purpose of this study was to determine the relative impact of small dam impoundments on the abundance of *Anopheles* mosquitoes in surrounding villages compared with smaller and more transient water bodies. Combining entomological surveillance with drone captured imagery and on-the-ground larval habitat surveys, the study focussed our efforts on Malawi’s dry season given the hypothesis that small dam impoundments have a greater impact on malaria during this period of the year. The study was conducted in Kasungu district which has recently been identified as a high priority district for malaria interventions in a recent malaria burden stratification exercise^17^. Furthermore, Kasungu reports a high percentage of its annual malaria cases during the dry season. In the year in which this study was conducted (2021), facilities in Kasungu reported a total of 431,277 confirmed malaria cases with 24% of these occurring in the dry months of June-October^18^.

## Methods

### Study area

This study was conducted in three communities in Kasungu district, Central Malawi which were situated approximately 7km from the district capital of Kasungu Town (Figure 1). These communities fall within UNICEF’s humanitarian drone testing corridor^19^ and form part of the *Maladrone* study which aims to (1) evaluate the feasibility of using drone imagery to identify mosquito larval habitat during the dry season, and (2) to use remotely sensed data on dry season larval habitat to measure the influence of habitat proximity on the spatial distribution of malaria risk. Further details relating to aim (1) can be found in Stanton et al. (2021)^20^, and further analysis using passive surveillance of malaria case data will be published elsewhere.

**Figure 1:**
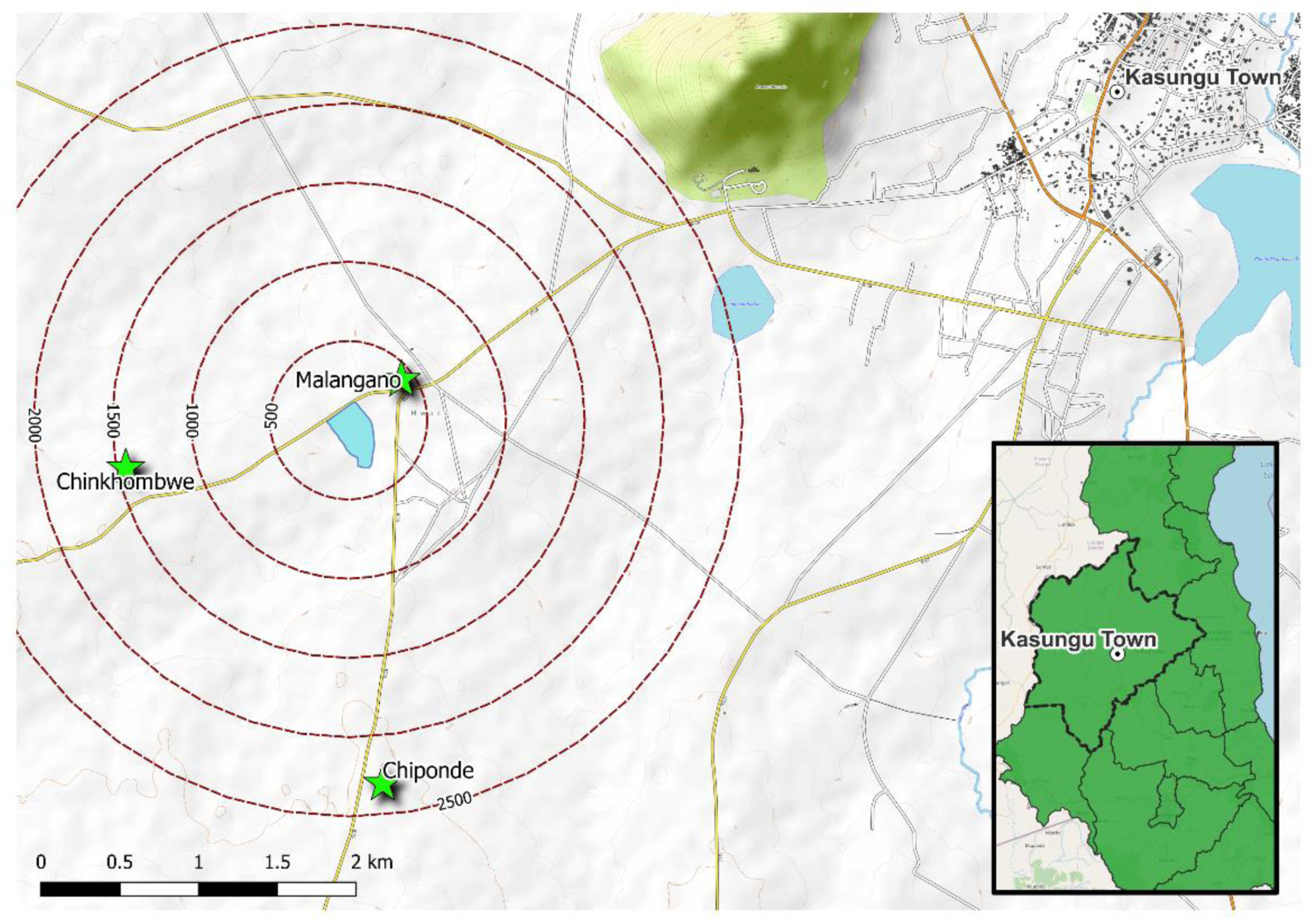
Map of the three study communities in relation to Kasungu town and surrounding dam impoundments.

The three communities of focus in this study were selected based on their proximity to a small dam which is being used by the local population to irrigate the surrounding area. The dam and its impoundment are situated next to Malangano village, covering an area of approximately seven hectares in the dry season (Figure 1). Previous larval surveys undertaken in the area confirmed the presence of *Anopheles* and c*ulicine* larvae within this impoundment during both the wet and dry season, indicating its likely influence on malaria transmission in the local area throughout the year. To determine the range over which the mosquito population is sustained by the impoundment during drier periods, in addition to undertaking mosquito surveys in Malangano village (total population of 165 households), two additional villages with houses situated approximately 1-1.5km (Chiponde, 92 households) and 1.8-2.6km (Chinkhombwe, 78 households) from the impoundment boundary were included in this study. An additional small dam was situated approximately 2km from houses in Malangano, with no additional small dams within 3.5km of the study communities. The nearest large dam was the Chitete dam, situated on the outskirts of Kasungu town.

### Household selection

A simulation study was conducted to determine the number of households to include in an indoor mosquito sampling exercise. Simulations were generated under the assumption that there needed to be a power of 80% to detect a range of differences in mosquito abundances (number of mosquitoes/trap/night) between the two communities separated by the greatest distance (Malangano and Chiponde). Based on the resources available to conduct this study, it was assumed that the smallest feasibly detectable difference was based on a relative risk of 0.5 i.e., to detect if the mosquito abundance in Chiponde was at most half of the abundance of Malangano. The selected study design comprised of the sampling of 30 households for two consecutive nights per month for four months, with the same households being sampled each month. Further details on the sample size simulation study are presented in Supplementary Materials A.

A list of households in each community was obtained from each village head, and 30 households were randomly selected using a random number generator. Consent to participate in the study was then sought from each of the selected household heads. If any household withdrew from the study during the four-month period, a replacement household was randomly selected from the household list.

Ethical approval for this study was obtained from both the Liverpool School of Tropical Medicine Research Ethics Committee (Protocol 20.004) and the Kamuzu University of Health Sciences Research Ethics Committee (Protocol P.07/19/2745). Further, a series of stakeholder engagement events were held to engage district officials and community leaders in the study, and community engagement activities took place within all three study communities prior to any recruitment taking place.

### Data collection

This study was conducted between May and August 2021, with data collection comprising of four components: household surveys, indoor collections of adult mosquitoes, drone image capture, and larval sampling.

#### Household surveys

Prior to starting the indoor mosquito collections, each household head was asked a series of questions to ascertain details about the household structure and composition including the number of people who usually slept in the house/the room where the trap was being placed, the number of bednets used in the house/room, roof materials and whether or not the eaves and windows were open or closed (Supplementary Materials B). Questionnaire data were captured on smartphones using OpenDataKit (ODK), with anonymised responses being uploaded and stored on Google Drive.

#### Indoor mosquito collections

CDC light traps (CDC-LT, Model 512; John W. Hock Company, Gainesville, Florida, USA) were placed overnight within the sleeping area of selected households to capture adult mosquitoes. Traps were set approximately 1.5m above the ground next to the foot of the bed between 6pm and 6am each trapping night. Households in each community were sampled sequentially over a two-week period, with approximately 15 houses being sampled each night. Collected mosquitoes were separated from other insects and were morphologically identified to genus level to separate the *Anopheles* from culicines. All mosquitoes were sexed as either males or females. Culicine mosquitoes were pooled together in storage bottles and no further identifications to species level were conducted. For the *Anopheles* mosquitoes, males were discarded while females were further identified as either *An. gambiae s*.l., *An. funestus s.*l and other anophelines. Individual female *Anopheles* were stored in 1.5 ml Eppendorf tubes containing a dessicant (silica gel) and each individual female *Anopheles* tubes were assigned unique IDs, and abdominal status (fed/unfed, gravid). All entomological data were subsequently recorded using ODK.

#### Drone image capture

Drone imagery was used to identify small water bodies in the study area which had the potential to harbour mosquito larvae. Imagery of the area surrounding each of the study villages was captured each month within the two-week period in which indoor mosquito collections were being conducted. Image capture was supported by nationally trained drone pilots via the drone service provider Globhe (https://www.globhe.com/). Due to the costs involved in capturing imagery it was necessary to limit the size of the area covered by the imagery. Images were therefore captured within a 400m buffer around the edges of each community only. Images were captured by multirotor drones (DJI Mavic Air and DJI Enterprise) and the resulting images were processed into orthomosaics with a spatial resolution of approximately 3cm. Due to the time required to upload the imagery and process the orthomosaic there was a time lag of approximately two weeks between the images being captured and the orthomosaic being available to the study team. Two members of the study team then independently reviewed each orthomosaic manually and each site where water was observed georeferenced and categorised. Categories included irrigation wells, small ponds, and irrigation channels. These two datasets were then reviewed together and consolidated, resulting in a single dataset for potential larval habitat locations for each community each month.

#### Larval sampling

Larval sampling was conducted during the same two-week window in which indoor adult mosquito collections were being undertaken. An adaptive sampling strategy was undertaken based on sampling data collected in the previous months, and practical considerations such as accessibility and weather. During the first sampling month, sampling took place along the shoreline of small dam impoundment at Malangano and in additional sites identified from ground-based surveys. The ground-based surveys were conducted with the support of local field staff and volunteers. In subsequent months, the previous month’s dataset of georeferenced potential larval habitat was used to guide the larval surveys. Sites were selected based on habitat category (irrigation wells, small ponds, pools around water pumps, dam impoundment) and accessibility. Due to time restrictions, some categories were excluded from the latter drier months’ surveys after no larvae were found in any of that category’s previously sampled sites in the early wetter months.

A standardised larval sampling approach was adopted for all habitat categories that were sampled. Larvae collections were done using sweep nets. Sweeping nets were dipped in the water at a 45- degree angle on the water surface, sweeping the entire accessible area (up to 1 metre wide) at once to minimize interruptions of the larvae. Sweeping was repeated 3 to 5 times, and all contents were emptied into a white plastic bowl and counted by genera (*Anopheles* or culicines) and by stage (early stage [L1/L2], late stage [L3/L4]). The number of pupae was also counted. Count data were recorded using ODK.

#### Additional data

Meteorological data were obtained for the study period to enable the variability in temperature and rainfall across the four-month period to be accounted for in subsequent statistical models of mosquito abundance. These data were obtained from ERA5-Land which combines ground data from meteorological stations with modelled data to produce hourly estimates at a 9km spatial resolution^21^. Data were extracted for the grid cells covering the study area, and aggregated measurements of total daily rainfall and minimum and maximum daily temperature were calculated from 1^st^ April 2021 up to the final data collection day (mid-August 2021).

### Data analysis

Summaries of the number of mosquitoes captured per household per sampling night were produced for each month, disaggregated by community, sex and genera. Further summaries of the female *Anopheles* were generated, including the number of each species complex identified by abdominal stage (unfed, blood-fed, half-gravid, gravid), disaggregated by community and month.

Focusing on female *Anopheles*, a univariate analysis of the community effect was undertaken. A simple generalised linear mixed model (GLMM) which included household-level random effects and a community effect only was fitted to the daily count data for each individual sampling month. A likelihood ratio test was then performed to compare this simple model to a null model (household-level random effects only) to assess whether significant differences between communities were present.

Summaries and plots of household characteristic data captured within household questionnaires were produced to explore the association between mosquito exposure risk, household composition (number of people who generally sleep in the room in which the net has been placed), the presence of bednets, and house construction (eaves open or closed, windows open or closed, roof type). Simple GLMMs fitted to the catch data aggregated over time were fitted which included household-level random effects and a single household characteristic fixed effects only. These models were compared to a base model which contained household-level random effects only via a likelihood ratio test.

Summaries of the types of potential larval habitat sites captured within the drone imagery were produced for each community by month. A 400m buffer was then created for each surveyed household using QGIS (version 3.28), and the number of potential sites identified in the buffer was calculated, in addition to the distance between the surveyed house and the boundary of the small dam impoundment at Malangano. Plots of these variables against mean female *Anopheles* catches per sampling night were generated to explore the association between mosquito exposure risk and surface water proximity, and simple GLMMs fitted to the monthly catch data were fitted which included household-level random effects and a surface water proximity fixed effect only. These models were again compared to a base model which contained household-level random effects only.

The number of sampled potential larval habitats, summarised by genus and stage (early/late), were tabulated by month and habitat type. Maps were produced of all identified potential larval habitat surrounding each community by month, with sampled habitats differentiated according to whether larvae were present or absent.

As weather conditions including temperature and rainfall impact the immature stages of the *Anopheles* lifecycle, temporally lagged effects of ERA5-Land derived variables on indoor adult mosquito catches were anticipated^22–24^. To explore this, the daily number of female *Anopheles* per trap per night was aggregated across all sampled houses, and the Pearson’s correlation between this value and lagged daily minimum, lagged maximum temperature, and lagged rainfall were calculated for lags of up to 21 days. To account for cumulative effects of temperature and rainfall conditions, moving averages were calculated with a window size of one and two weeks, such that 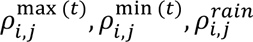 represent the correlation between weather conditions lagged by *i* days, averaged over *j* = 1,7,14 days for maximum daily temperature, minimum daily temperature and total daily rainfall respectively. As such, conditions up to 35 days prior to the trapping date were taken into consideration. Values of *i* and *j* which maximised the correlation were reported, and simple univariate GLMMs were fitted to the daily data and compared to a base model using the corresponding lagged and averaged weather variables.

More complex spatio-temporal GLMMs were fitted to the mosquito count data to further quantify the influence of household-level characteristics and the surrounding environment on indoor catches. Counts were initially assumed to follow a Poisson distribution, and the fitted GLMMs were tested for overdispersion. Three sets of models were fitted. In the initial model (Model 1), household characteristics were considered as fixed effects to explain between-household differences in catch numbers. Community name and month were included as fixed effects to crudely account for spatial and seasonal differences in catch data respectively. Random effects in Model 1 included collection date to account for differences in localised condition affecting indoor catches and household ID to account for unmeasured differences in households (Supplementary Materials C). To explore whether seasonal differences could be fully or partially explained by the weather conditions, Model 2 incorporated additional fixed effects using variables derived from the ERA5-Land dataset. Finally, Model 3 explored whether any further spatio-temporal variability could be explained by the presence or absence of small potential larval habitats within 400m of the house. Akaike’s Information Criterion (AIC) was calculated for each of the models under consideration within each model set, and the results of the model with the lowest AIC were reported. The marginal and conditional R^2^ values, calculated using methods reported in Nakagawa et al. 2017^26^, were generated for each of the selected models to evaluate the variance explained by the fixed effects in the model and the fixed and random effects combined. Model diagnostics were performed on the best overall model with respect to AIC which included tests for overdispersion, heteroscedasicity, collinearity and the normality of random effects.

All data analyses were conducted in R (v4.3.0), with GLMMs fitted using the `glmer` function within the `lme4` (version 1.1-29) package. Model diagnostics were generated using the `check_model` function within the `performance` package (version 0.9.0)^25^.

## Results

Table 1 displays the number of mosquitoes caught and mean catch per trap per night by genus and sex in each of the three study communities for each sampling month (May-August 2021). In total, 1,256 mosquitoes were captured, of which 32.8% (N = 412) were female *Anopheles*, 58.8% (N = 739) were female culicines, 1.7% (N = 21) were male *Anopheles* and 6.7% (N = 84) were male culicines. Catches were consistently higher in Malangano across all sampling months, with the largest differences observed in June and July 2021. Figure 2 presents the mean number of female *Anopheles* and culicines per trap per day for each community over time and their respective 95% confidence intervals. Simple GLMMs fitted to the catch data for each individual month indicate that in May, at the start of the dry season, there were no significant difference (p=0.2343) in mean female *Anopheles* catches between the three communities (1.54 [95% CI: 0.926, 2.154], 0.96 [95% CI: 0.439,1.481] and 1.29 [95% CI: 0.797, 1.783] per trapping day for Malangano, Chinkhombwe and Chiponde respectively). While catches declined over time in all three communities, the decline was sharpest in the two communities further from the dam, whereas catches in Malangano remained significantly higher (p<0.0001) in both June (1.37 [95% CI: 0.820, 1.920]) and July (0.85 [95% CI: 0.489, 1.211]). In August, a significant difference remained (p=0.0087), although mean catches in all three communities were low. The lowest catches in August were observed in Chiponde (0.10 [95% CI: 0.013, 0.187]), with Malangano and Chinkhombwe being marginally higher (0.42 [95% CI: 0.220, 0.620], 0.25 [95% CI: 0.137,0.363] per trapping day for Malangano and Chinkhombwe respectively). For culicines, a similar pattern was observed for May (p=0.1263), June (p<0.0001) and July (p<0.0001), however in August (p=0.0023) catches remained relatively high in Malangano (1.47 [95% CI: 0.350, 2.590]) and Chiponde (0.93 [95% CI: 0.359, 1.501]) compared to Chinkhombwe (0.15 [95% CI: 0.033, 0.267]).

**Figure 2:**
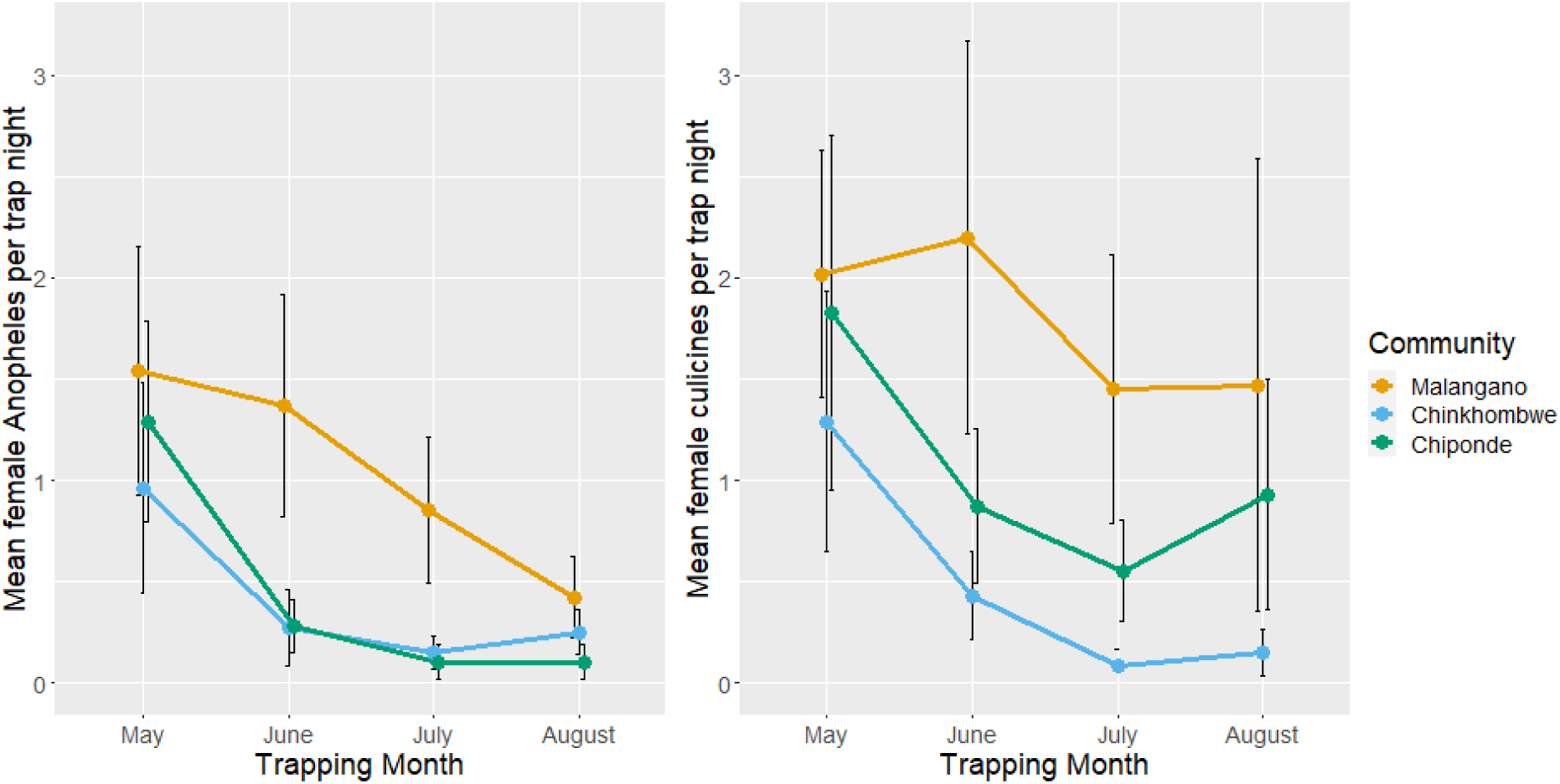
Mean number of female *Anopheles* (left) and culicine (right) mosquitoes captured in each community by month. Bars represent the 95% confidence intervals.

**Figure 3:**
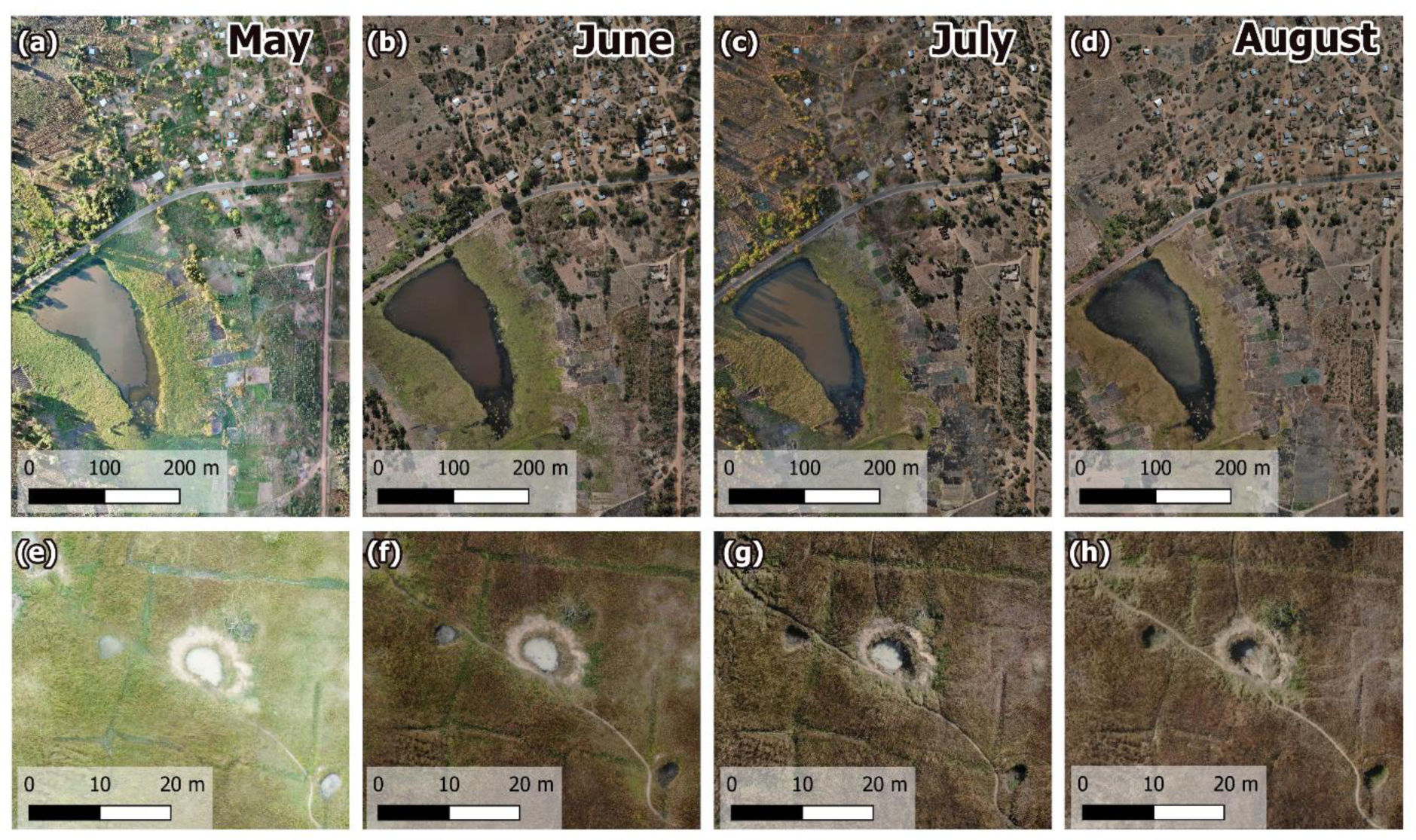
Drone imagery of the study area for May-August 2021. (a)-(d) display the small dam impoundment, with irrigation well and channels shown in (e)-(h).

**Table 1:**
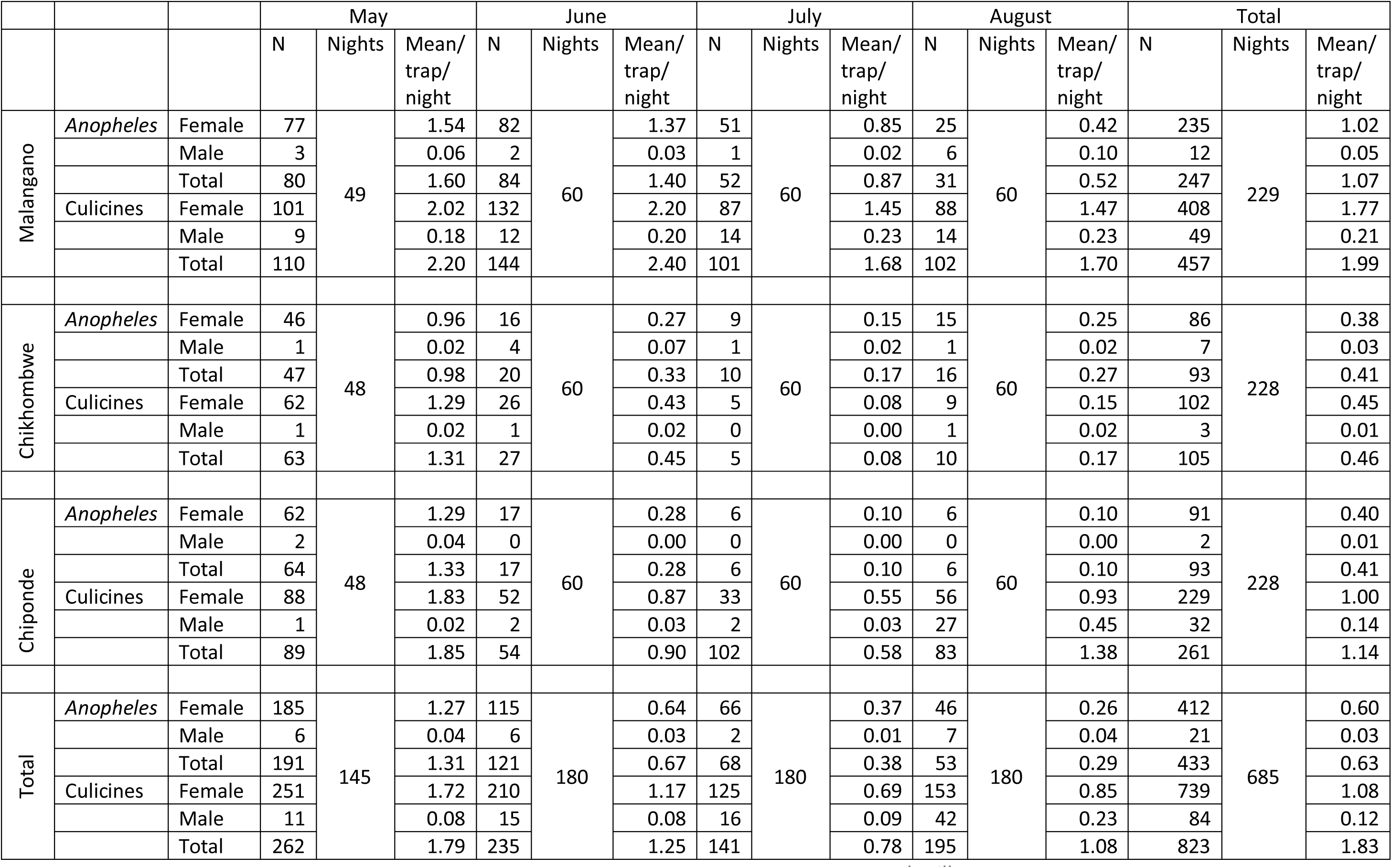
Summaries of mosquito trap data by genera, sex and month for houses within the three study villages.

|  |  |  | May |  |  | June |  |  | July |  |  | August |  |  | Total |  |  |
| --- | --- | --- | --- | --- | --- | --- | --- | --- | --- | --- | --- | --- | --- | --- | --- | --- | --- |
|  |  |  | N | Nights | Mean/<br>trap/<br>night | N | Nights | Mean/<br>trap/<br>night | N | Nights | Mean/<br>trap/<br>night | N | Nights | Mean/<br>trap/<br>night | N | Nights | Mean/<br>trap/<br>night |
| Malangano | *Anopheles* | Female | 77 | 49 | 1.54 | 82 | 60 | 1.37 | 51 | 60 | 0.85 | 25 | 60 | 0.42 | 235 | 229 | 1.02 |
|  |  | Male | 3 |  | 0.06 | 2 |  | 0.03 | 1 |  | 0.02 | 6 |  | 0.10 | 12 |  | 0.05 |
|  |  | Total | 80 |  | 1.60 | 84 |  | 1.40 | 52 |  | 0.87 | 31 |  | 0.52 | 247 |  | 1.07 |
|  | *Culicines* | Female | 101 |  | 2.02 | 132 |  | 2.20 | 87 |  | 1.45 | 88 |  | 1.47 | 408 |  | 1.77 |
|  |  | Male | 9 |  | 0.18 | 12 |  | 0.20 | 14 |  | 0.23 | 14 |  | 0.23 | 49 |  | 0.21 |
|  |  | Total | 110 |  | 2.20 | 144 |  | 2.40 | 101 |  | 1.68 | 102 |  | 1.70 | 457 |  | 1.99 |
| Chikhombwe | *Anopheles* | Female | 46 | 48 | 0.96 | 16 | 60 | 0.27 | 9 | 60 | 0.15 | 15 | 60 | 0.25 | 86 | 228 | 0.38 |
|  |  | Male | 1 |  | 0.02 | 4 |  | 0.07 | 1 |  | 0.02 | 1 |  | 0.02 | 7 |  | 0.03 |
|  |  | Total | 47 |  | 0.98 | 20 |  | 0.33 | 10 |  | 0.17 | 16 |  | 0.27 | 93 |  | 0.41 |
|  | *Culicines* | Female | 62 |  | 1.29 | 26 |  | 0.43 | 5 |  | 0.08 | 9 |  | 0.15 | 102 |  | 0.45 |
|  |  | Male | 1 |  | 0.02 | 1 |  | 0.02 | 0 |  | 0.00 | 1 |  | 0.02 | 3 |  | 0.01 |
|  |  | Total | 63 |  | 1.31 | 27 |  | 0.45 | 5 |  | 0.08 | 10 |  | 0.17 | 105 |  | 0.46 |
| Chiponde | *Anopheles* | Female | 62 | 48 | 1.29 | 17 | 60 | 0.28 | 6 | 60 | 0.10 | 6 | 60 | 0.10 | 91 | 228 | 0.40 |
|  |  | Male | 2 |  | 0.04 | 0 |  | 0.00 | 0 |  | 0.00 | 0 |  | 0.00 | 2 |  | 0.01 |
|  |  | Total | 64 |  | 1.33 | 17 |  | 0.28 | 6 |  | 0.10 | 6 |  | 0.10 | 93 |  | 0.41 |
|  | *Culicines* | Female | 88 |  | 1.83 | 52 |  | 0.87 | 33 |  | 0.55 | 56 |  | 0.93 | 229 |  | 1.00 |
|  |  | Male | 1 |  | 0.02 | 2 |  | 0.03 | 2 |  | 0.03 | 27 |  | 0.45 | 32 |  | 0.14 |
|  |  | Total | 89 |  | 1.85 | 54 |  | 0.90 | 102 |  | 0.58 | 83 |  | 1.38 | 261 |  | 1.14 |
| Total | *Anopheles* | Female | 185 | 145 | 1.27 | 115 | 180 | 0.64 | 66 | 180 | 0.37 | 46 | 180 | 0.26 | 412 | 685 | 0.60 |
|  |  | Male | 6 |  | 0.04 | 6 |  | 0.03 | 2 |  | 0.01 | 7 |  | 0.04 | 21 |  | 0.03 |
|  |  | Total | 191 |  | 1.31 | 121 |  | 0.67 | 68 |  | 0.38 | 53 |  | 0.29 | 433 |  | 0.63 |
|  | *Culicines* | Female | 251 |  | 1.72 | 210 |  | 1.17 | 125 |  | 0.69 | 153 |  | 0.85 | 739 |  | 1.08 |
|  |  | Male | 11 |  | 0.08 | 15 |  | 0.08 | 16 |  | 0.09 | 42 |  | 0.23 | 84 |  | 0.12 |
|  |  | Total | 262 |  | 1.79 | 235 |  | 1.25 | 141 |  | 0.78 | 195 |  | 1.08 | 823 |  | 1.83 |

Of the 412 female *Anopheles* captured during the indoor trapping exercise, 91% (373/412) were morphologically identified as *An. funestus sl.*, 4% (18/412) were *An. gambiae s.l*, and 5% (21/412) were other secondary anopheline species (unidentified) (Table 2). Most of the *An. gambiae s.l.* mosquitoes were captured within houses in Malangano in the first month of sampling (11/18), and none were captured in July or August in any of the sampled houses. A total of 52 (13%) of the 412 female *Anopheles* were blood fed (28 blood fed, 22 gravid, 2 semi-gravid), with the majority of these (41) being captured in Malangano, five in Chikhombwe and six in Chiponde (Supplementary Materials D).

**Table 2:** Females *Anopheles* captured in each community by month, summarised by morphologically identified species complex.

|  |  | May | June | July | August | Total |
| --- | --- | --- | --- | --- | --- | --- |
| Malangano | *An. funestus* | 57 | 77 | 49 | 24 | 207 |
|  | *An. gambiae* | 11 | 2 | 0 | 0 | 13 |
|  | Other | 9 | 3 | 2 | 1 | 15 |
|  | Total | 77 | 82 | 51 | 25 | 235 |
| Chinkhombwe | *An. funestus* | 44 | 15 | 7 | 14 | 81 |
|  | *An. gambiae* | 1 | 1 | 0 | 0 | 2 |
|  | Other | 1 | 0 | 2 | 1 | 4 |
|  | Total | 46 | 16 | 9 | 15 | 86 |
| Chiponde | *An. funestus* | 60 | 15 | 6 | 5 | 86 |
|  | *An. gambiae* | 2 | 1 | 0 | 0 | 3 |
|  | Other | 0 | 1 | 0 | 1 | 2 |
|  | Total | 62 | 17 | 6 | 6 | 91 |
| Total | *An. funestus* | 161 | 107 | 62 | 43 | 373 |
|  | *An. gambiae* | 14 | 4 | 0 | 0 | 18 |
|  | Other | 10 | 4 | 4 | 3 | 21 |
|  | Total | 185 | 115 | 66 | 46 | 412 |

Household characteristics of the sampled houses are summarised in Table 3 by community. Communities were similar with respect to the number of residents in each household (median of 4 in Malangano and Chikhombwe, 4.5 in Chiponde), and number of people sleeping in the room in which the CDC-LT was situated (2 in Malangano and Chikhombwe, 3 in Chiponde). Bed net ownership was highest in Chiponde (all 30 houses owned at least one net), and lowest in Malagano (21/31 owned at least one net). With regards to house construction, there were no clear differences between communities with approximately half of surveyed houses having an iron sheet roof and half having a thatched roof in all three communities. Only three houses had tiled roofs (2 in Chinkhombwe, 1 in Chiponde). The eaves were open or partially open in approximately half of surveyed houses in all three communities, whereas most houses (76/92, 83%) had open or partially open windows.

**Table 3:** Household characteristics by study community. *Open includes partially open.

|  |  | Malagano | Chinkhombwe | Chiponde |
| --- | --- | --- | --- | --- |
| N sampled houses |  | 31 | 31 | 30 |
| Median residents | $\leq 5$ | 1 | 1 | 1 |
| | $> 5$ | 3 | 4 | 3 |
|  | Total | 4 | 4 | 4.5 |
| Median no. people sleeping in room with trap | $\leq 5$ | 0 | 0 | 0 |
| | $> 5$ | 2 | 2 | 2 |
|  | Total | 2 | 2 | 3 |
| N. houses with at least one net |  | 21 | 25 | 30 |
| Mean nets per resident |  | 0.30 | 0.28 | 0.39 |
| N. houses with at |  | 20 | 23 | 29 |
| least one net in room with trap |  |  |  |  |
| Mean nets per person in room with trap |  | 0.30 | 0.38 | 0.44 |
| Roof type | Iron sheets | 15 | 15 | 14 |
|  | Thatched | 16 | 14 | 15 |
|  | Tiles | 0 | 2 | 1 |
| Eaves | Closed | 14 | 15 | 14 |
|  | Open* | 17 | 16 | 16 |
| Windows | Closed | 7 | 5 | 4 |
|  | Open* | 24 | 26 | 26 |

Whilst household characteristics may not explain differences in mosquito catches between communities, their influences on within-community variability of aggregated catches of female *Anopheles* were explored (Supplementary Materials E). These figures, combined with the results of the simple GLMMs, indicated that there were no significant differences in females *Anopheles* per trap per sampling night for all sampling months combined and the number of people usually sleeping in the room (p=0.4405), whether or not there was at least one bednet in the room (p=0.5321), and whether or not windows were closed or (partially) open (p=0.8728). Significantly higher catches were however observed between houses with thatched roofs compared to iron roofs (p=0.0221) and between houses with (partially) open eaves compared to closed eaves (0.0171). It was noted that most houses with iron roofs had closed eaves (38/44), and thatched roofs had open eaves (39/45).

Drone imagery captured for each of the four study months clearly shows a decline in green vegetation over the period in all areas excluding the periphery of the impoundment next to Malangano village (Figure 4). Manual reviews of the imagery, independently undertaken by two team members, identified additional small water bodies in the vicinity of the three communities, primarily in the form of small irrigation wells (Figure 4), the majority of which were in the surrounds of Chiponde (28 observed in May, compared to 8 surrounding Malagano and 11 surrounding Chinkhombwe). Chikhombwe had far fewer potential larval habitat sites in its immediate surroundings compared to the other two communities. Most sites dried up over the observation period in Chinkhombwe and Chiponde (Table 4), whereas the number of irrigation wells in Malangano was sustained as a result of several new wells being created close to the impoundment in July and August.

**Figure 4:**
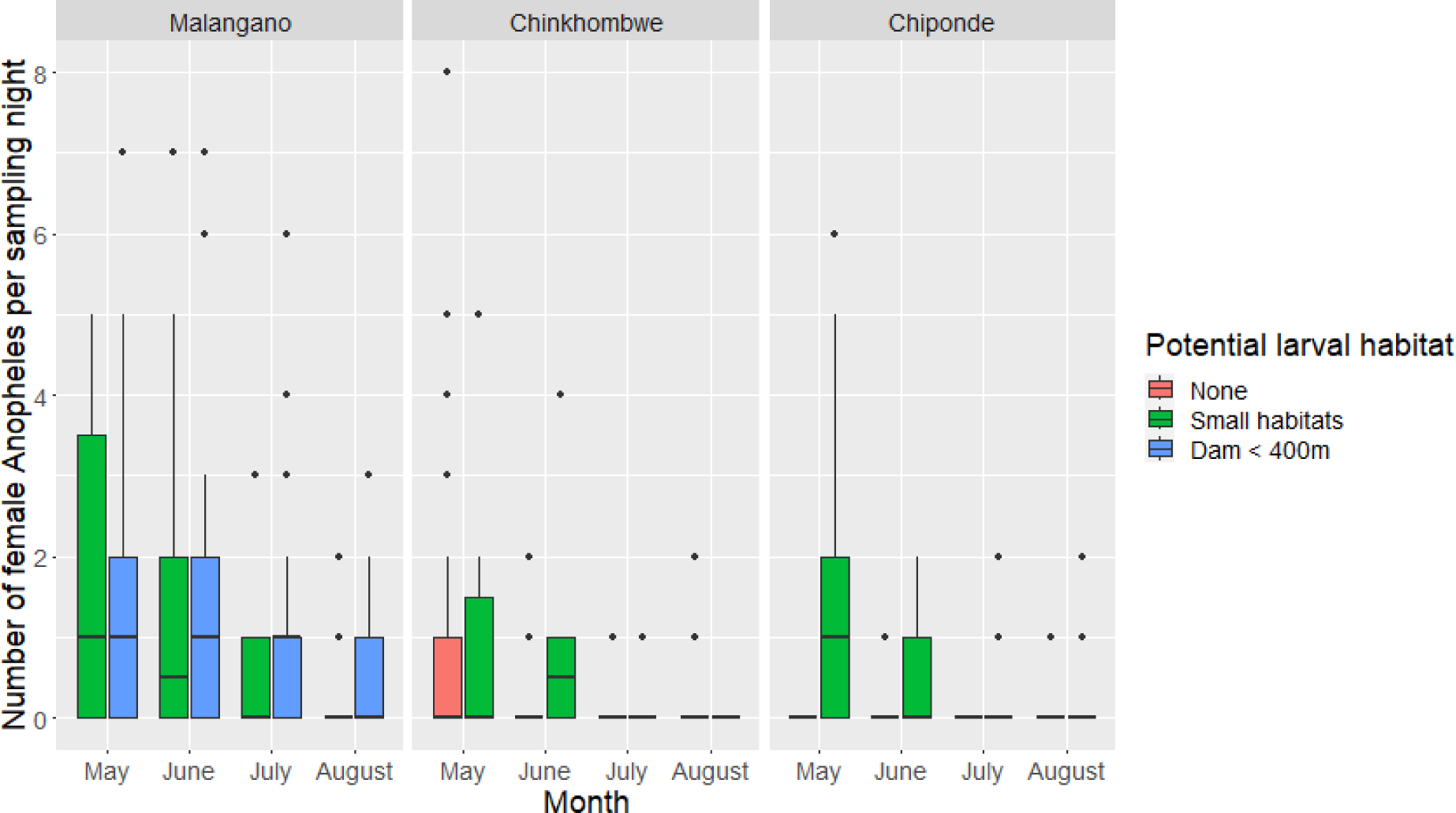
Number of female *Anopheles* catches per sampling night by potential larval habitat proximity for each study month by study community.

**Table 4:** Number of water bodies detected through manual examination of drone imagery within the surrounds (400m) of households included in the study.

|  |  | May | June | July | August |
| --- | --- | --- | --- | --- | --- |
| Malangano | Irrigation wells | 8 | 5 | 7 | 7 |
|  | Ponds | 6 | 4 | 0 | 0 |
|  | Pooling around water pumps | 5 | 4 | 2 | 2 |
|  | Puddles/irrigation channels | 3 | 1 | 1 | 0 |
|  | Total | 22 | 14 | 10 | 9 |
| Chikhombwe | Irrigation wells | 11 | 5 | 4 | 2 |
|  | Ponds | 0 | 0 | 0 | 0 |
|  | Pooling around water pumps | 0 | 0 | 0 | 0 |
|  | Puddles/irrigation channels | 4 | 0 | 0 | 0 |
|  | Total | 15 | 5 | 4 | 2 |
| Chiponde | Irrigation wells | 28 | 21 | 9 | 4 |
|  | Ponds | 3 | 2 | 0 | 0 |
|  | Pooling around water pumps | 1 | 1 | 1 | 1 |
|  | Puddles/irrigation channels | 3 | 1 | 0 | 0 |
|  | Total | 32 | 28 | 10 | 5 |

Figure 4 presents boxplots of female *Anopheles* catches per night by potential larval habitat proximity categories (no habitats within 400m, only small habitats within 400m, Malangano’s dam within 400m) by sampling month. The presence of small potential habitats (irrigation wells etc.) near houses in Chikhombwe and Chiponde appear to have higher indoor catches than those houses with no potential habitat nearby for May and June only, with this effect diminishing in July and August. Simple GLMMs fitted to the household-level data for each month indicate a significant difference in female *Anopheles* catches in June (0.0006), July (<0.0001) and August (0.0103). Given the higher number of small habitats in the vicinity of Chiponde in comparison to Chikhombwe in May and June (Table 4), this may explain why a greater mosquito abundance was observed in Chiponde over this period despite it being situated the furthest from Malangano’s dam.

To explore the potential for these small water bodies as larval habitats, preliminary larval sampling of 13 sites within the study area was undertaken in May 2021 using local field staff’s knowledge of the area. In June-August, additional sampling sites were selected by combining information extracted from the drone imagery with local knowledge. A summary of sites sampled by habitat type is presented in Table 5. Due to the one-month delay between the drone imagery and larval sampling activities, an additional 23 drone-identified sites were visited by the field team (10 in June, 6 in July, 7 in August) that had since dried up and are therefore not included in this summary.

**Table 5:** Number of sites sampled for larvae by habitat type and month.

|  | May | June | July | August | Total |
| --- | --- | --- | --- | --- | --- |
| Irrigation wells | 7 | 4 | 10 | 16 | 37 |
| Ponds | 2 | 7 | 2 | 0 | 11 |
| Pooling around water pumps | 2 | 2 | 5 | 0 | 9 |
| Impoundment | 2 | 1 | 5 | 7 | 15 |
| Total | 13 | 14 | 22 | 23 | 72 |

Summaries of the results of the larval sampling by month and habitat type are presented in Table 7. Additional summaries by month and stage (early or late) are presented in Supplementary Materials F. Due to the nature of the sample site selection, it is not possible to use these data to draw any robust conclusions about trends over time for each habitat type. Irrigation wells, for which the most samples are available, indicate that while the proportion of sites where no larvae were observed is high (60%, 25/37), they plausibly contribute to malaria transmission in dryer months. Maps of the sampled sites for each month in relation to the sampled households are presented in Figure 5.

**Figure 5:**
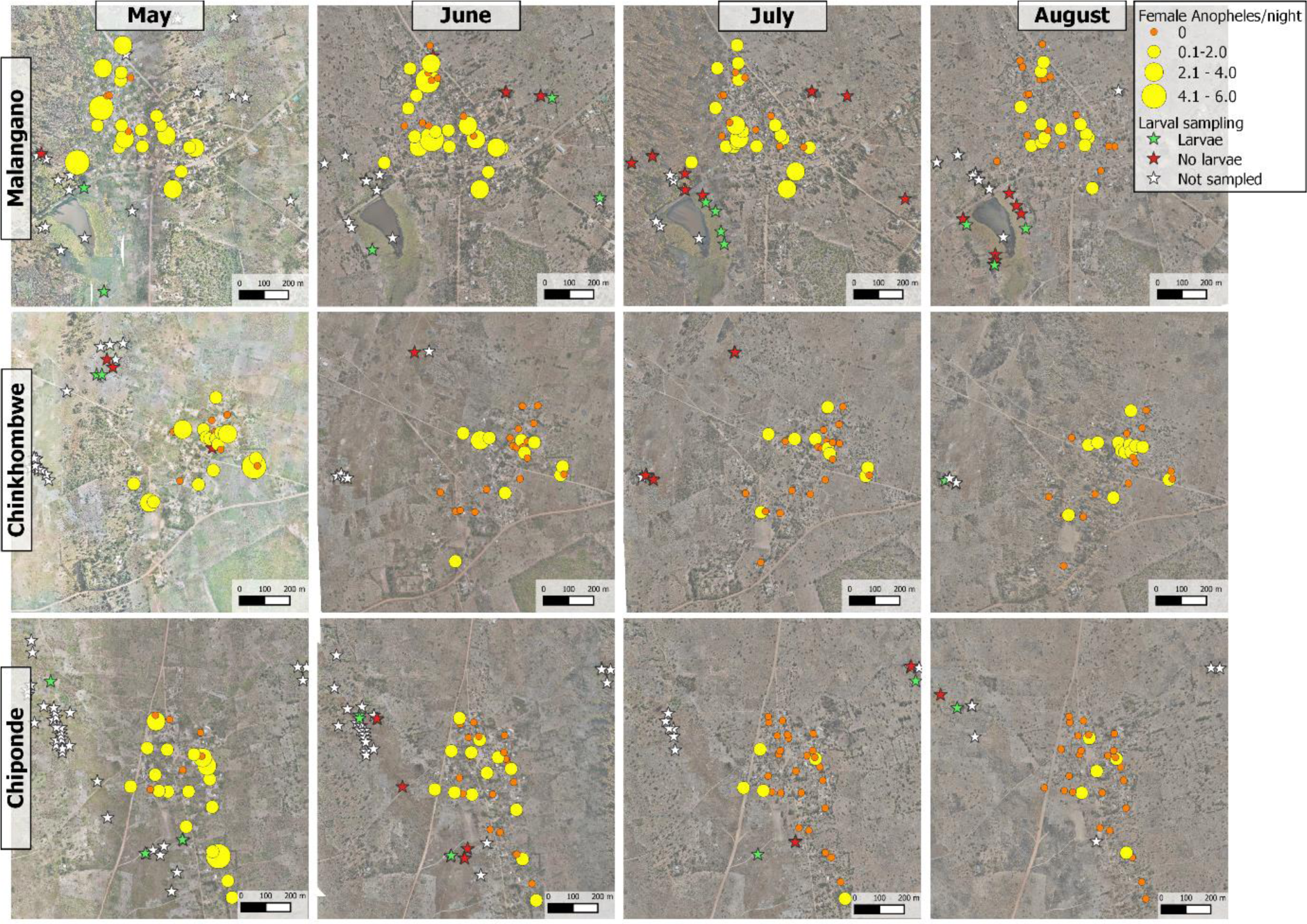
Entomological data consisting of household-level number of female *Anopheles* per trap per night and larval sampling sites by larval presence/absence for each community by month.

**Table 7:**
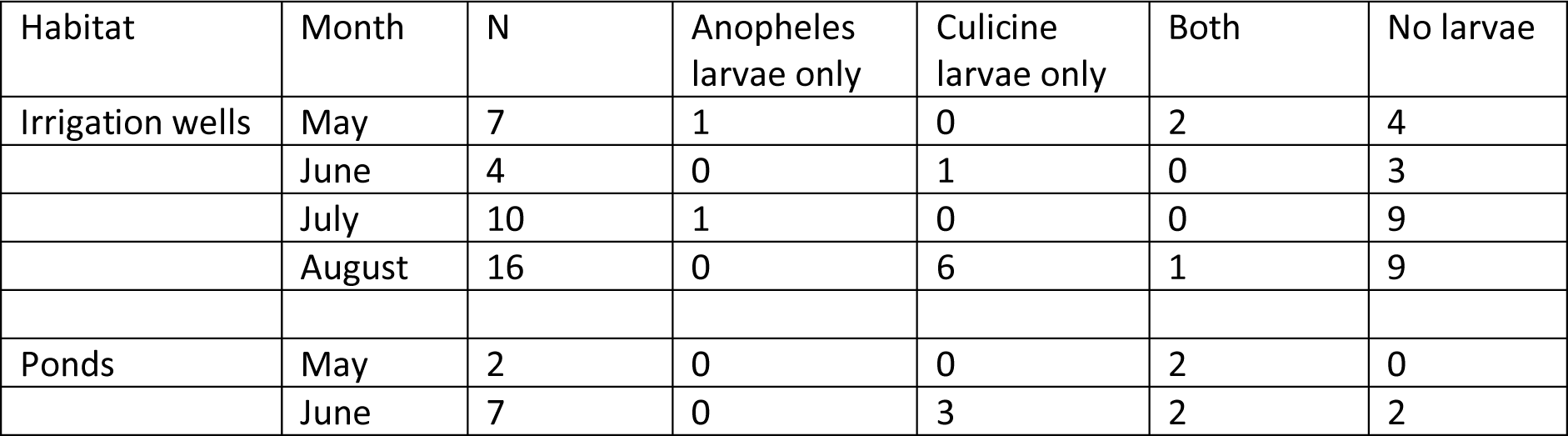
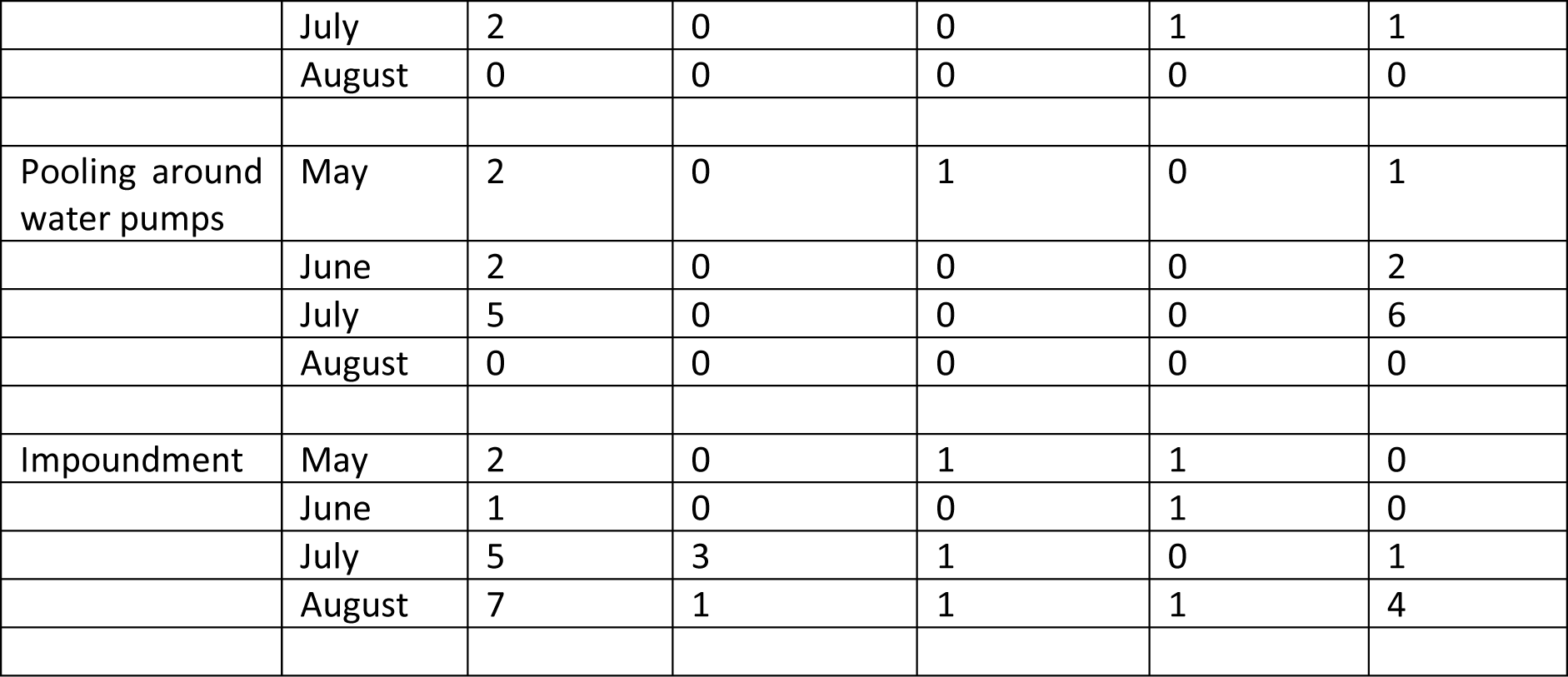
Larval sampling data by habitat type and month.

ERA5-Land weather data comprising of hourly temperature and rainfall were extracted for the 9km-by-9km grid cell containing the study area for April to August 2021 inclusive using the `ncdf4` package (version 1.21) in R. Hourly data were aggregated to daily summary statistics (minimum temperature, maximum temperature and total rainfall) (Figure 6). In the month prior to the start of the indoor surveys, daily minimum temperature ranged between 13 to 18 °C (median = 16 °C), with temperatures lowering to 8-16 °C (median 11 °C) over the 4-month study period, rising again after the final collections had been conducted. Daily maximum temperature was less variable, ranging between 22-27 °C (median = 24 °C) in the month prior to the study and lowering slightly to 20 – 26 °C (median = 23 °C) during the study. In the month prior to the study collections starting, daily rainfall ranged between 0 and 7 mm (median = 0.4 mm), whereas during the study period there was very little rainfall (min = 0, max = 1, median = 0), with just 31 of the 103 days (30%) between the first and last collections recording more than 0.01 mm. To account for the lagged and cumulative effects of temperature and rainfall on mosquito development, the Pearson’s correlation (*ρ*_*i*,*j*_) between temperature lagged by *i* = 1, . ., 21 days and averaged over *j* = 1,7, 14 days against the daily female *Anopheles* catch data were generated (Figure 7). Generally, a stronger correlation was observed when averaging over the previous 14 days compared to considering the weather on a single day, with a small amount of variation across the considered time lags. A stronger correlation was observed between daily female *Anopheles* catches and minimum temperature (max(*ρ*_*i*,*j*_) = *ρ*_20,14_ = 0.75) and total rainfall (max(*ρ*_*i*,*j*_) = *ρ*_20,14_ = 0.68), compared to maximum temperature (max(*ρ*_*i*,*j*_) = *ρ*_10,14_ = 0.56). Simple univariate GLMMs fitted to the optimal weather variables such that the weather variable was included as a linear fixed effect and the collection date was included as a random effect indicated a significant positive relationship (p<0.001) between all three weather variables and daily female *Anopheles*/trap/night.

**Figure 6:**
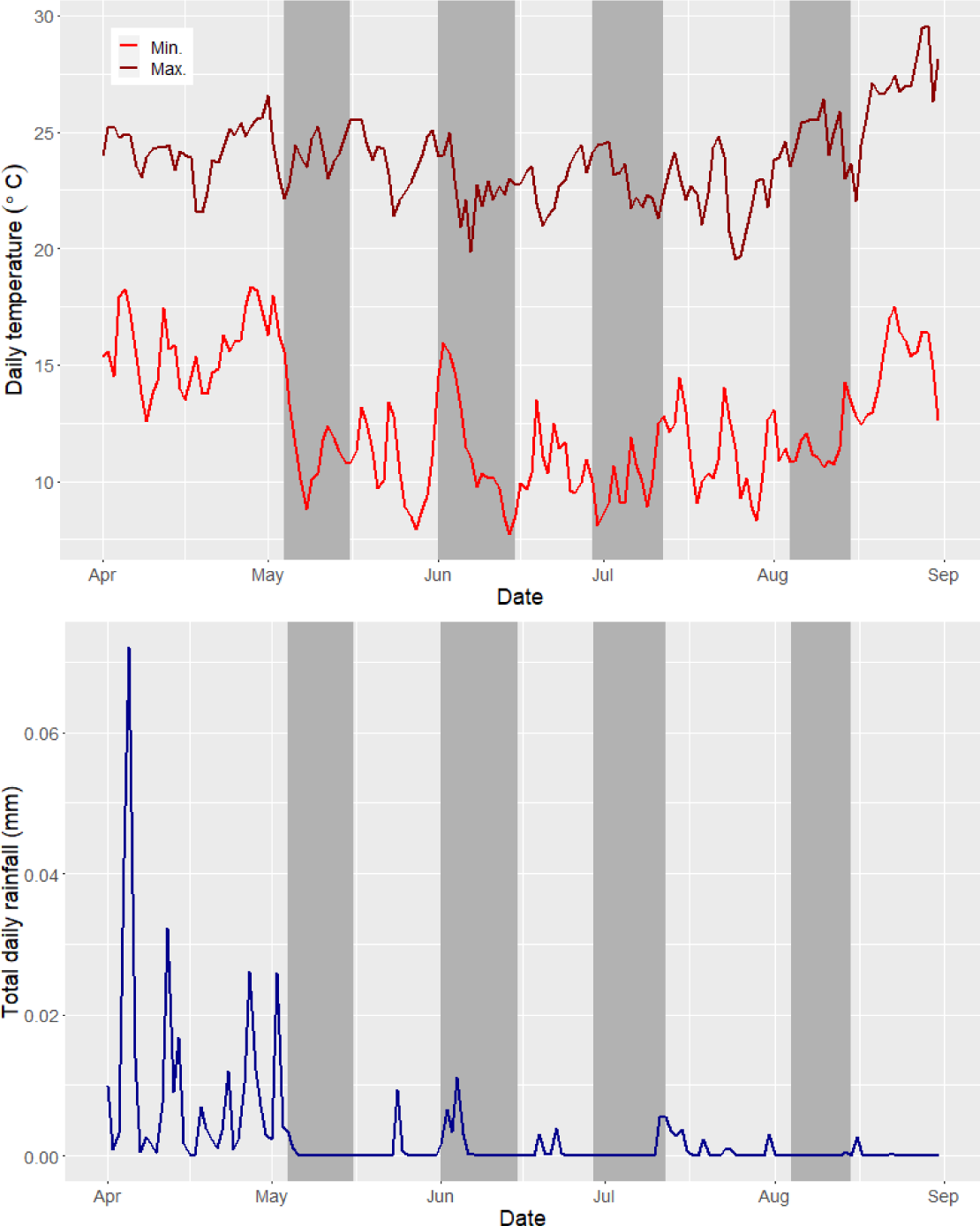
Daily minimum and maximum temperature and total rainfall for the study area, derived from ERA5-Land data. Dark grey shaded sections indicate indoor mosquito sampling periods.

**Figure 7:**
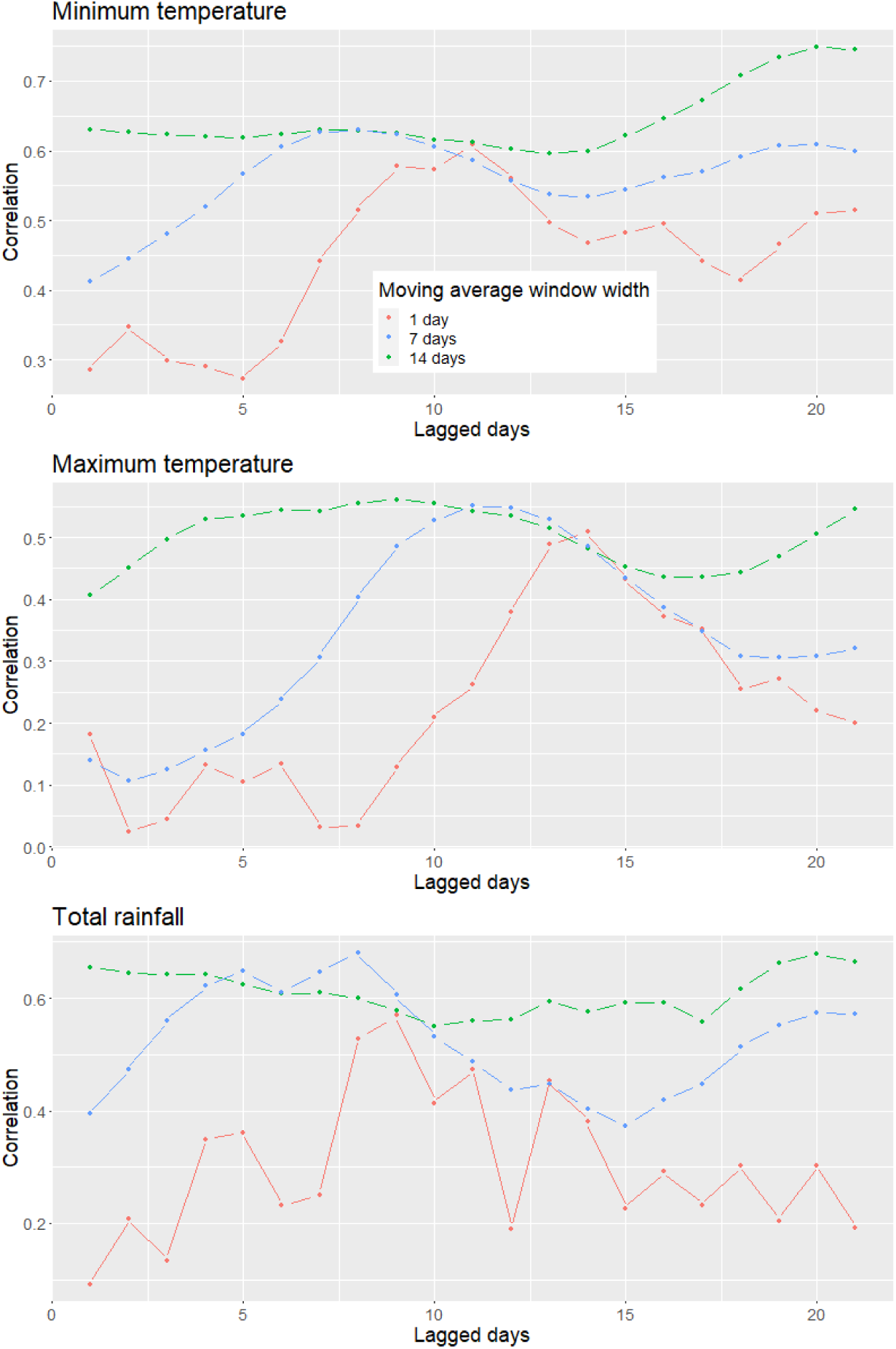
Pearson’s correlation between daily number of female *Anopheles*/trap/night and moving averages (1 [red line], 7 [blue line]and 14 [green line] days) lagged by 1-21 days.

Building upon the above exploratory analysis, a series of GLMMs were fitted to the daily household-level female *Anopheles* abundance data (Table 8). Initially, community name and collection month were included as fixed effects to crudely capture spatial and temporal differences, with household-level and community-level random effects being incorporated to account for further unmeasured differences (Model 1, AIC = 1293, marginal R^2^ = 0.29, conditional R^2^ = 0.50). Compared to baseline (catches in Malagano in May in houses with closed eaves) it was observed after adjusting for other fixed effects, catches were significantly smaller in Chinkhombwe (relative risk [RR] = 0.32 [95% CI 0.18 – 0.55]) and Chiponde (RR = 0.31 [95% 0.18 – 0.54]). Catches declined significantly in successive months (June RR = 0.43 [95% CI 0.28-0.66], July RR = 0.25 [95% CI 0.16 – 0.40], August RR = 0.18 [0.11 – 0.30]). Catches were significantly higher in houses with (partially) open eaves compared to closed eaves (RR = 1.93 [95% CI 1.33 – 2.78]).

**Table 8:**
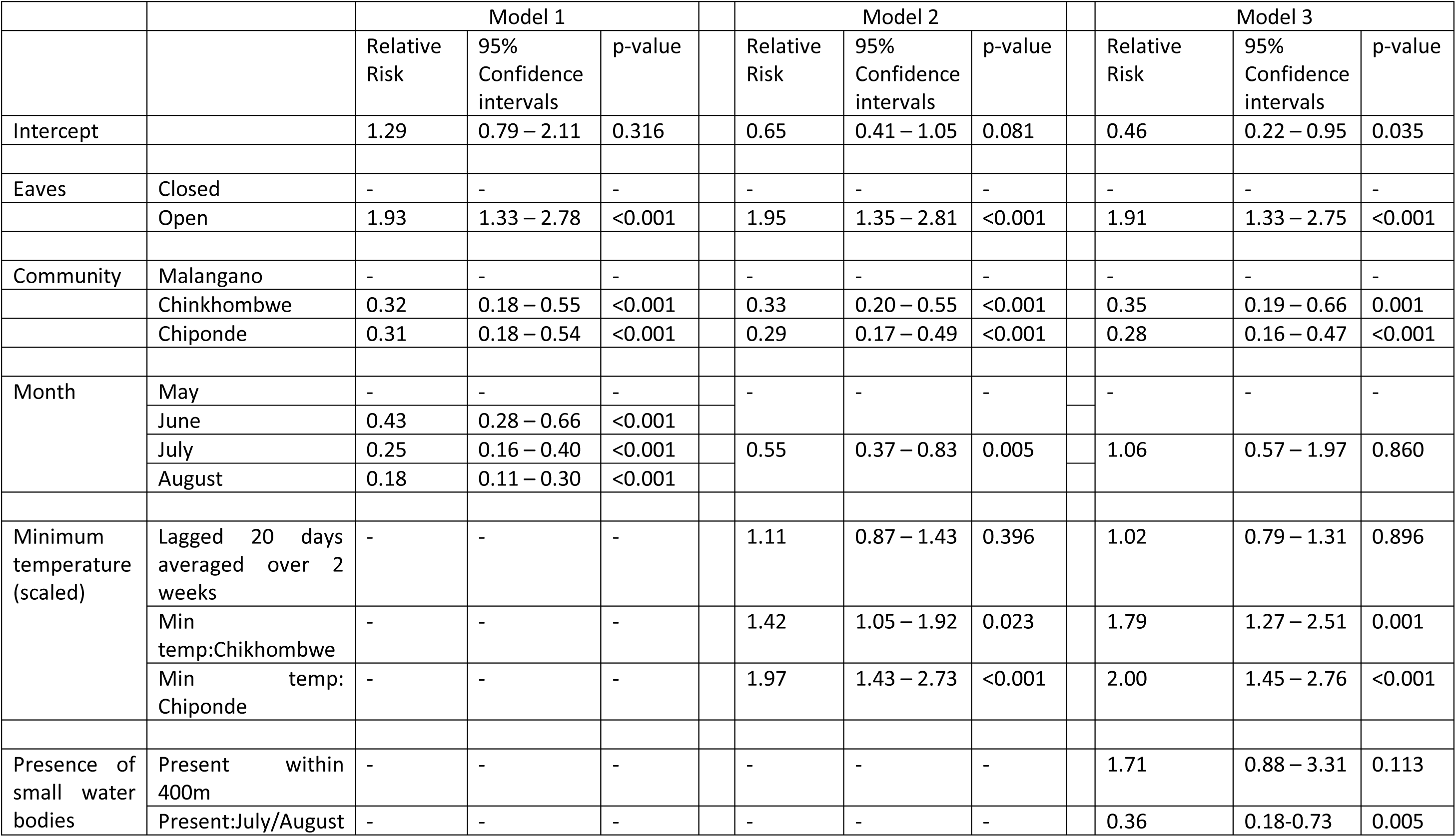
Fixed effects for best fitting GLMM within each model set - Models 1 (AIC = 1293, Marginal R^2^ = 0.29), Model 2 (AIC = 1280, Marginal R^2^ = 0.34), Model 3 (AIC = 1275, Marginal R^2^ = 0.35).

Weather variables in the form of minimum temperature averaged over two weeks, lagged by 20 days, maximum temperature averaged over two weeks lagged by 10 days, and total rainfall summed over two weeks, lagged by 20 days were considered in Model 2 to account for temporal variability in female *Anopheles* catches. A community effect was included in the model to account for both crude spatial variability and as a proxy for proximity to the Malangano dam. Exploratory plots of weather against catches indicated that the relationship varied between the three study communities, which is likely due to the influence of the dam (Supplementary Materials G). An interaction between community and weather was therefore considered to account for this variability. Using AIC for model comparison, the minimum temperature variable best captures the variability in catches over time (Table 8). Model 2 had an AIC of 1280, with a marginal R^2^ of 0.34 and a conditional R^2^ of 0.51. The effect of minimum temperature on catches varied between communities. In Malangano (the baseline community), no significant relationship was observed (RR = 1.11, [95% CI 0.87 – 1.43]), whereas a positive significant relationship was demonstrated in Chinkhombwe (interaction RR = 1.42, [95% CI 1.05 – 1.92]) and Chiponde (interaction RR = 1.97 [95% CI 1.43 – 2.73]). The inclusion of minimum temperature explains some of the variability in catches over time, however after adjusting for minimum temperature changes, the catches were still lower than expected in July/August compared to May/June (IRR = 0.55, [95% CI 0.37 – 0.83]). Note that time was grouped into two-month periods in Model 2 as opposed to single months as it was observed that the grouping of months resulted in a marginally lower AIC and the simpler model was therefore favoured.

The inclusion of proximity to larval habitat was incorporated into Model 3 (AIC = 1275, marginal R^2^ = 0.35, conditional R^2^ of 0.51). Due to the design of the study, proximity to the dam was already captured in the community effect. Proximity to small water bodies was included as a binary variable (small water present/absent within 400m). The number of small water bodies within 400m was considered, however the inclusion of this additional information did not significantly improve the model. An interaction between the presence of water and month was included to account for the observation that catches were very low in July/August regardless of water being detected in the area. In May/June (baseline time period), a non-significant increase in relative risk was observed in houses with small water bodies identified within 400m (RR = 1.71 [95% CI 0.79 – 1.31]). This effect was however significantly reduced in July/August (interaction RR = 0.36 [95% CI 0.18 – 0.73]).

Model diagnostics were performed on the best fitting model (Model 3). No significant overdispersion was detected (p=0.494). Further, no substantial evidence of heteroscedasicity was detected, and distributional assumptions about the random effects in the model appeared reasonable (Supplementary Materials H). A boxplot of observed against fitted values (Supplementary Materials I) indicate that the model captures the general trend in catches, differentiating reasonably well between low and high catches, but generally underpredicts larger observed values.

## Discussion

*Anopheles* mosquito abundance during the dry season was highly focal within our study, with the focality seemingly driven by the presence of a small dam and its impoundment. The influence of proximity to this impoundment on indoor mosquito abundance dropped much more sharply than anticipated, with significantly fewer *Anopheles* caught in villages situated just 1 - 2.6 km from the impoundment periphery. This extended impact on mosquito numbers, however, lasted only two months into the dry season with *Anopheles* abundances falling considerably thereafter. The relative contribution of smaller water bodies on *Anopheles* production in the area - identified from drone-captured imagery and on-the-ground surveys - was inconclusive but any contribution certainly diminished into the dry season. Most of the female *Anopheles* caught by indoor CDC light traps were *An. funestus*, suggesting that small dam impoundments are colonised by this predominant vector of malaria during the dry season in Kasungu.

Water harvesting structures, whether large power generating hydrodams or smaller community-based dams, perpetuate malaria transmission in Africa^27^. Most of the epidemiological and entomological evidence gathered to date has focussed on larger dam structures^8^ but meta-analyses across the continent suggest that smaller dams may have an even greater role to play^28^. In an analysis of malaria incidence in populations living less than 5 km from the shoreline to small and large dams in four river basins across Africa, smaller dams contributed to 77-85% of cases^28^. Here, our primary goal was to determine how temporal changes in the local landscape surrounding a small dam in the dry season – in an area designated as ‘highest malaria burden’ by the NMCP – influences mosquito abundance and diversity. Proximity to the dam had a clear influence on indoor female *Anopheles* catch numbers in the adjacent village of Malangano. Shortly after the rains subsided in May, *Anopheles* densities were similar in all three study villages, but as the dry season continued into June and July, households in Malangano received significantly higher numbers of female *Anopheles* compared with households in Chiponde and Chikhombwe; villages situated at maximal distances of 1.5 km and 2.6 km from the dam respectively. While small differences were observed in bed net ownership and housing structure between communities, large between community differences were still observed after adjusting for this within the GLMM framework. Furthermore, whereas there is a positive significant relationship between lagged minimum temperature and *Anopheles* abundance observed in Chiponde and Chikhombwe, no such relationship is observed in Malangano, indicative of an overriding effect of the dam on mosquito numbers in this village.

The perimeter of small dam impoundments provide excellent habitat for the larvae of *Anopheles* malaria vectors^7^. Small dams are generally earthen structures relying on natural drainage with shallow edges often supporting a variety of floating and/or emergent vegetation. Furthermore, receding waters in the dry season leave exposed soil and clay that create transient pools. This creates a wider variety of mosquito habitat around the periphery compared with larger heavily constructed dams. In one of the few detailed larval ecology studies focussed on small dams^7^, vegetation and soil type along the shoreline had a significant impact on *Anopheles* species abundance and richness, with associations between habitat type and species corresponding to known ecologies and population dynamics of the vector. Small dam impoundments are not homogenous habitats and extensive heterogeneity in mosquito habitat can exist around a single dam.

Small dams are scattered across the district of Kasungu providing a vital source of water for domestic and agricultural activities. We observed nearly zero rainfall during the period of mosquito collections (May to August) and so these dam impoundments represent one of the few permanent habitats available for *Anopheles* to lay their eggs during the dry season. This is supported by drone-captured declines in green vegetation across the area and the disappearance of smaller alternative water bodies such as irrigation wells and ponds by the end of August, presumably through evapotranspiration (Fig. 3). Earlier on in the dry season, these smaller water bodies do appear to influence *Anopheles* abundance in households situated within 400m, particularly in Chiponde and Chikhombwe (Figure 4) but overall, their relative contribution remains equivocal compared to the dam impoundment.

*An. funestus* was the primary *Anopheles* vector collected in households suggesting that small dam impoundments act as dry season habitat for this vector. The relative importance of *An. funestus* in malaria transmission in Malawi has increased over the past decade^29^ echoing similar species shifts across East Africa^30,31^ due to its high anthropophilic behaviour and efficiency as a vector of *P. falciparum*. In national annual vector monitoring surveys using CDC-LTs during the same period of our study (2021), *An. funestus* was the predominant vector in two sites sampled within Kasungu District^29^. Seasonal entomological collections in different parts of Tanzania have shown that populations and entomological rates of *An. funestus* are higher in the dry season^30,32^, consistent with its preference for permanent water bodies. These breeding sites of *An. funestus* have been, however, notoriously difficult to find compared with *An. gambiae s.l.* but recent extensive ecological surveys in Tanzania show that this vector prefers to lay eggs in open semi-permanent water bodies with emerging vegetation and clear water as well as in those habitats exposed to sunlight and high temperatures^33^ – fitting the description of the type of small dam impoundments found across Kasungu, including the one in our study. While we did not perform any species level larval identification, the lack of any other semi-permanent water bodies in Kasungu, strongly suggests that small dam impoundments support *An. funestus* populations in Malawi. The small percentage of *An. gambiae s.l.* were all captured by June, suggesting their presence was associated with rainfall at the end of the wet season.

The CDC-LT data suggest that the vast majority of *An. funestus* emerging as adults from the small dam impoundment flew a short distance (<500m) to households in Malangano. Mosquito dispersal is influenced by many factors including odour plume^34^, host availability and density^35^, wind patterns and the spatial configuration of households and villages^36^. The sharp decline in *An. funestus* abundance in Chiponde and Chikhombwe in July and August suggests that the dam and village of Malangano is acting as a ‘hotspot’ for *An. funestus*, with the presence of human settlements next to the dam essentially limiting mosquito dispersal and concentrating vector densities. Whether this interaction between mosquito breeding site and the spatial configuration of houses intensifies or extends malaria transmission into the dry season within our setting is unknown. Human biting rates are expected to occur closest to the breeding site^37^ whereas exposure to infectious, older mosquitoes, should occur further away from the site of adult emergence^38^. Indeed, the relative risk of catching indoor *Anopheles* in households with open eaves (confirming previous findings in Malawi^39^) demonstrates the importance of other factors such as house construction.

What are the practical implications of our findings? Our investigation into the impact of small dams on mosquito populations and malaria transmission aims to inform recommendations for additional vector control that complement frontline tools like insecticide-treated nets. Based on our findings, small dam impoundments provide focal habitat for the most efficient malaria vector in Malawi, and targeting these areas with larval source management (LSM) could have substantial benefits on those communities living within their vicinity. The optimal timing and coverage of LSM is determined by the local vector ecology and seasonality of malaria, balanced with the practicalities of implementation. The current WHO recommendation is that LSM is most impactful in the dry season, yet a recent modelling study using the OpenMalaria platform suggests that LSM is more effective at the beginning or during the rainy season in seasonal settings^40^. Investment in LSM as part of an integrated vector management strategy remains subject to debate and is context specific, but the small dams situated in Kasungu are worthy of exploring LSM as a viable option. A community-based approach employing a range of environmental management activities, significantly reduced both larval and adult *An. arabiensis* densities around a small dam in highland Ethiopia where malaria transmission is markedly seasonal^41^.

To conclude, this study highlights the influence of human-made structures, specifically small dams, on malaria vector exposure in Central Malawi. As these small dams continue to play a vital role in food security and climate change resilience across sub-Saharan Africa, urgent measures are needed to mitigate their public health impact. The highly localised effects of dam impoundments on vector densities revealed in this study can inform the design of targeted control methods, including LSM. Future research should prioritise cost-effective strategies for implementing these targeted interventions.

## Acknowledgements

We would like to acknowledge the Maladrone fieldworkers, Geoffrey Chaundwa and Blessings Phiri and community volunteers John Banda, Odeta Chiwanga and Vincent Lisutha for the valuable support they provided during the indoor mosquito trapping and larval sampling activities. We would further like to thank the residents of Malangano, Chikhhombwe and Chiponde, particularly those who allowed us to place traps in their homes, for welcoming us into their communities and supporting this study.

## Author contribution

**Kennedy Zembere**: Investigation (lead); resources (lead); project administration (equal); supervision (supporting); writing – original draft preparation (supporting). **Christopher M Jones**: Conceptualisation (equal); funding acquisition (supporting); methodology (equal); supervision (equal); writing – original draft preparation (equal). **Rhosheen Mthawanji:** Investigation (supporting); resources (supporting); writing – review and editing (supporting). **Clinton Nkolokosa:** Investigation (supporting); resources (supporting); writing – review and editing (supporting). **Richard Kamwezi:** Investigation (supporting); resources (supporting); writing – review and editing (supporting). **Patrick Ken Kalonde:** Investigation (supporting); resources (supporting); writing – review and editing (supporting). **Michelle C Stanton:** Conceptualisation (equal); data curation; formal analysis; funding acquisition (lead); methodology (equal); project administration (equal); supervision (equal); visualisation, writing – original draft preparation (equal).

This article includes a reflexivity statement detailing the authors’ approach to equitable international collaboration.

## Data availability statement

The household-level entomological data that support the findings of this study are openly available in Zenodo at https://doi.org/10.5281/zenodo.8414747.

Household information data are openly available in Zenodo at https://doi.org/10.5281/zenodo.10067093. Household coordinates have been removed from this dataset due to privacy restrictions.

Drone imagery relating to this study are openly available from openaerialmap.org.

## Ethical approval statement

This study was approved by the Kamuzu University of Health Sciences ethics committee (Protocol number P.07/19/2745) and Liverpool School of Tropical Medicine Research Ethics committee (Protocol number 20-004).

