## Supplementary Materials for "Small dams drive *Anopheles* abundance during the dry season in a high malaria burden area of Malawi"

A. Sample size simulation study

To determine the number of houses from which to sample mosquitoes, a sample size calculation was undertaken based on ensuring that there will be enough power to detect a difference between the mosquito abundance in the community to the reservoir (Malangano) in comparison to the nearest community to Malangano (Chinkhombwe). Sample size simulation were undertaken under a range of scenarios assuming the abundance data are realisations from the following generalised linear mixed model.

Let $Y_{ijk}$ be the random variable representing the number of mosquitoes caught in one trap over a single night of sampling. We assume $y_{ijk} \sim Pois(\lambda_{ijk})$ where *i* represents the house, *j* represents the community, and *k* represents the sampling day. Then $\log(\lambda_{ijk}) = \eta_{ijk}$ represents the linear predictor such that

$$\eta_{ijk} = \beta_0 + \sum_{j=1}^{2} \beta_j z_{ij} + u_j + \nu_{ij} + \epsilon_{ijk}$$

where $\beta_0$ is the average number of expected mosquitoes per house per day on the log scale in Malangano, $z_{ij}$ is an indicator of the distance band in which the house is situated, with j=1 to 2. $\beta_j$ is the associated effect of distance band on the number of mosquitoes per house per day relative to the reference distance band (log-relative risk). The $u_j$ represents the community-level random effects which we assume to have a normal distribution with variance $\gamma^2$and $\nu_{ij}$ represents house-level variation which is nested within villages, with variance $\tau^2$ and $\epsilon_{ijm}$ represents the observation-level random effect which we include to account for over-dispersion. We assume this has a normal distribution with variance $\sigma^2$.

We perform a simulation study to determine how many households we would need to sample to achieve a power of 80% to detect a difference between the number of mosquitoes caught in Malangano in comparison to Chinkhombwe. The smallest difference we wish to detect is based on a relative risk of 0.5 i.e. we want to be able to detect if the number of mosquitoes caught in Chinhombwe is at most half of the number caught in Malangano. In addition to considering the number of houses we also consider the impact of varying values of $\beta_0$, $\gamma^2$, $\nu_{ij}$, $\sigma^2$ in addition to the number of nights over which each household is sampled. Using the `sim.glmm` function within `the GLMMmisc` package in R, we conducted 1000 simulations for each scenario being considered. Figure A presents the results of one set of simulations where it was assumed that the number of sampled houses per village ranged from 40 to 60. The estimated power for each scenario is plotted, assuming village-level variance ($\gamma^2$) of 0.1 and 0.5, and overdispersion variance ($\sigma^2$) of 0.1 and 0.5. The plots are similar when considering sampling each house in one village for two or three nights, ranging house-level variance ($\tau^2$) between 0.1 and 0.5.

Based on these results we sampled 30 houses per distance band over two nights, allowing for 1-2 dropouts per village.

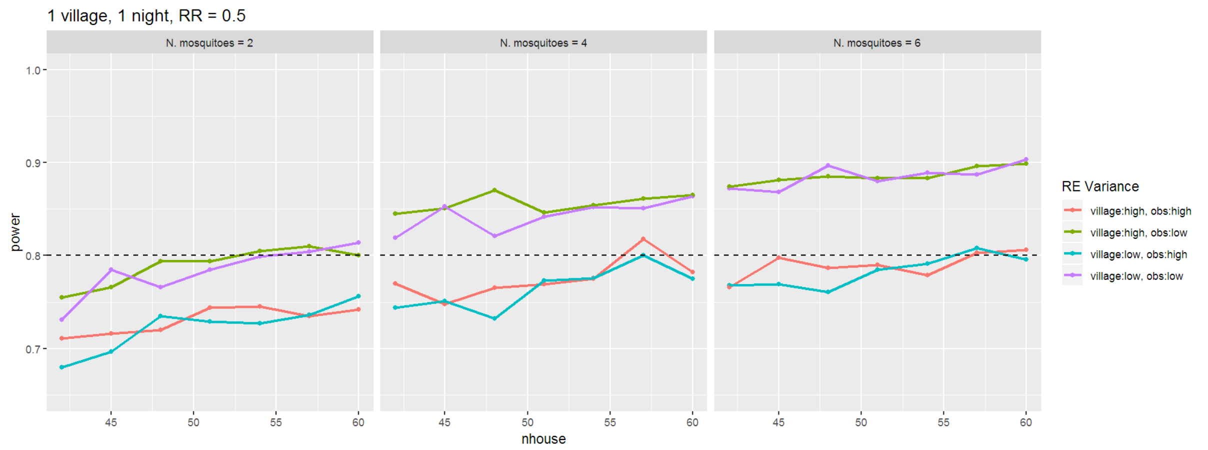

**Figure A: Results of the sample size simulations assuming one village per distance band is sampled, with each house being sampled for one night.**

B. Household questionnaire

Each household head responded to the following questions prior to the first deployment of the CDC light traps within their household.

- Number of people in the household, disaggregated by age (aged over 5 and aged under 5)?
- Number of people who usually sleep in the room in which the trap will be situated (aged over 5 and aged under 5)?
- Number of bednets used by the household?
- Number of bednets used in the room in which the trap will be situated?
- Number of people who usually sleep under a bednet in the household (under 5 and over 5)?
- Number of people who usually sleep under a bednet in the room where the trap will be situated (under 5 and over 5)?

The research team also noted the following details about the construction of the house:

- Roof type (thatched, iron sheets or tiles)
- Status of eaves (closed, partially open, open)
- Status of windows (closed, partially open, open, none)

C. Model specification

Let $Y_{ijk}$ be the random variable representing the number of mosquitoes caught in one trap situated in household $i$, community $j$ on sampling night $k$. We assume $y_{ijk} \sim Pois(\lambda_{ijk})$, and $\log(\lambda_{ijk}) = \eta_{ijk}$ represents the linear predictor. We consider three approaches to modelling $\eta_{ijk}$.

Model 1: Covariates include household characteristic $x_{ij}$, community ($C_j$) and month of collection ($M_k$). Random effects include a household-level random intercept ($U_i$) and a collection night random intercept ($V_k$) which are both assumed to have a normal distribution with zero mean and variance $\tau^2$ and $\theta^2$ respectively.

$$\eta_{ijk} = \alpha + \boldsymbol{\beta}' x_{ij} + \gamma C_j + \delta M_k + U_i + V_k$$

Model 2: In addition to the covariates included in Model 1, this model also incorporates weather variables to determine whether they can be used to explain any temporal variation observed in the data. Each weather variable $w_{\mathrm{k-l,n}}$ is lagged by $l$ days and averaged over the previous $n$ days, n=1,7 or 14. As an example, $w_{k-6,7}$ represents the weather variable averaged over days $k-12$ to $k-6$ (7 days).

$$\eta_{ijk} = \alpha + \boldsymbol{\beta}' x_{ij} + \gamma C_j + \delta M_k + \theta w_{l-k,n} + U_i + V_k$$

Model 3: This model explored whether any further spatio-temporal variability could be explained by the presence or absence of small potential larval habitats within 400m of the house. Using the locations of water bodies identified from drone imagery, an indicator variable $I_{ijk}$ was derived such that $I_{ik} = 1$ if a water body was identified within 400m of household $i$ in community $j$ in month $M_k$, and zero otherwise.

$$\eta_{ijk} = \alpha + \boldsymbol{\beta}' x_{ij} + \gamma C_j + \delta M_k + \theta w_{l-k,n} + \tau I_{ijk} + U_i + V_k$$

D. Status of female anophelines by sampling month and community

| Community | | May | June | July | August | Total |
|---|---|---|---|---|---|---|
| Malangano | Bloodfed | 5 | 8 | 4 | 4 | 21 |
| | Gravid/semi-gravid | 0 | 8 | 10 | 2 | 20 |
| | Non-gravid | 72 | 66 | 37 | 19 | 194 |
| | Total | 77 | 82 | 51 | 25 | 235 |
| Chinkhombwe | Bloodfed | 1 | 0 | 0 | 0 | 1 |
| | Gravid/semi-gravid | 3 | 0 | 0 | 1 | 4 |
| | Non-gravid | 42 | 16 | 9 | 14 | 81 |
| | Total | 46 | 16 | 9 | 15 | 86 |
| Chiponde | Bloodfed | 2 | 3 | 1 | 0 | 6 |
| | Gravid/semi-gravid | 0 | 0 | 0 | 0 | 0 |
| | Non-gravid | 60 | 14 | 5 | 6 | 85 |
| | Total | 62 | 17 | 6 | 6 | 91 |
| Total | Bloodfed | 8 | 11 | 5 | 4 | 28 |
| | Gravid/semi-gravid | 3 | 8 | 10 | 3 | 24 |
| | Non-gravid | 174 | 96 | 51 | 39 | 360 |
| | Total | 185 | 115 | 66 | 46 | 412 |

E. Boxplots of number of female *Anopheles* per trap per night by household characteristics.

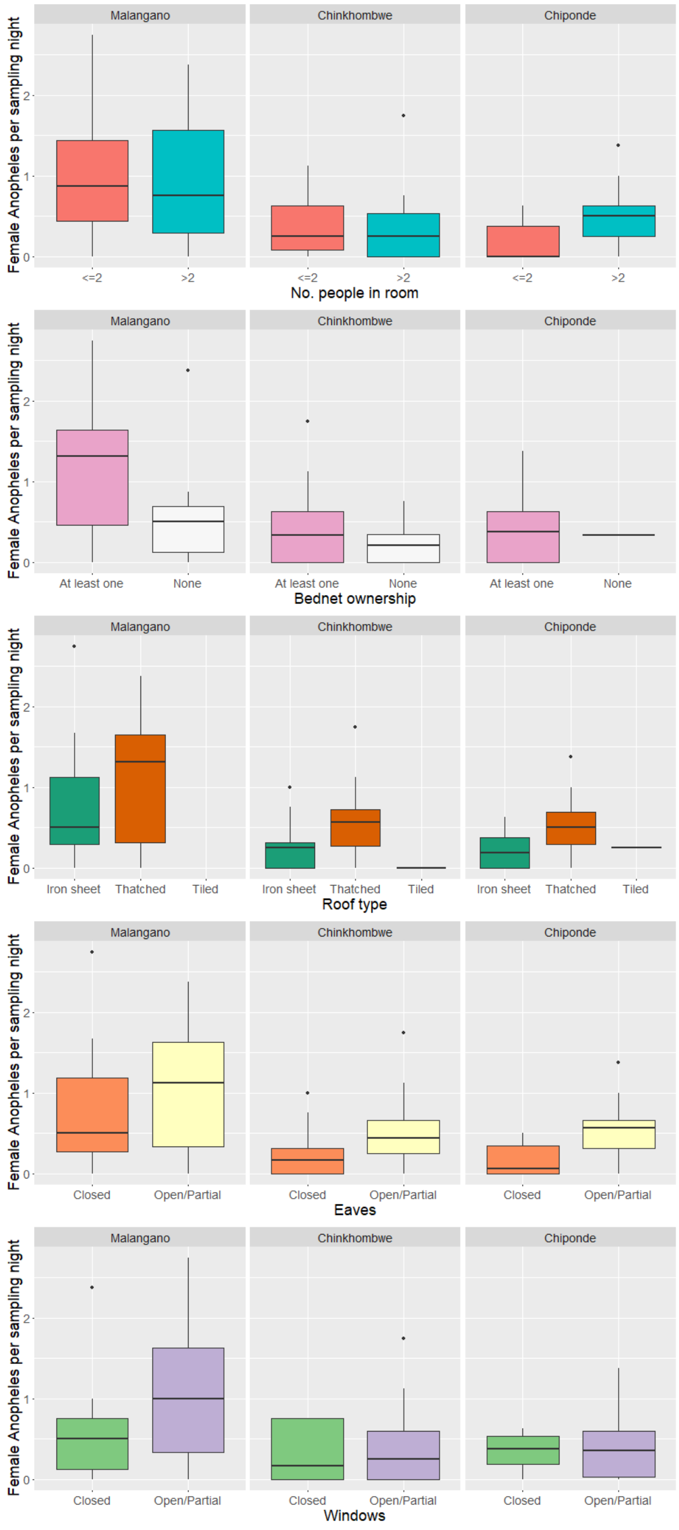

F. Larval sampling results by genera, stage and month.

| | | May | June | July | August |
|---|---|---|---|---|---|
| N sites | | 13 | 14 | 22 | 23 |
| *Anopheles* | Early stage | 6 | 3 | 5 | 3 |
| | Late stage | 2 | 1 | 1 | 1 |
| Culicines | Early stage | 7 | 6 | 2 | 8 |
| | Late stage | 3 | 5 | 1 | 3 |
| None (%) | | 5 (38%) | 7 (50%) | 16 (73%) | 13 (57%) |

G. Plots of abundance against averaged weather conditions. Solid line represents smoothed conditional means generated using LOESS.

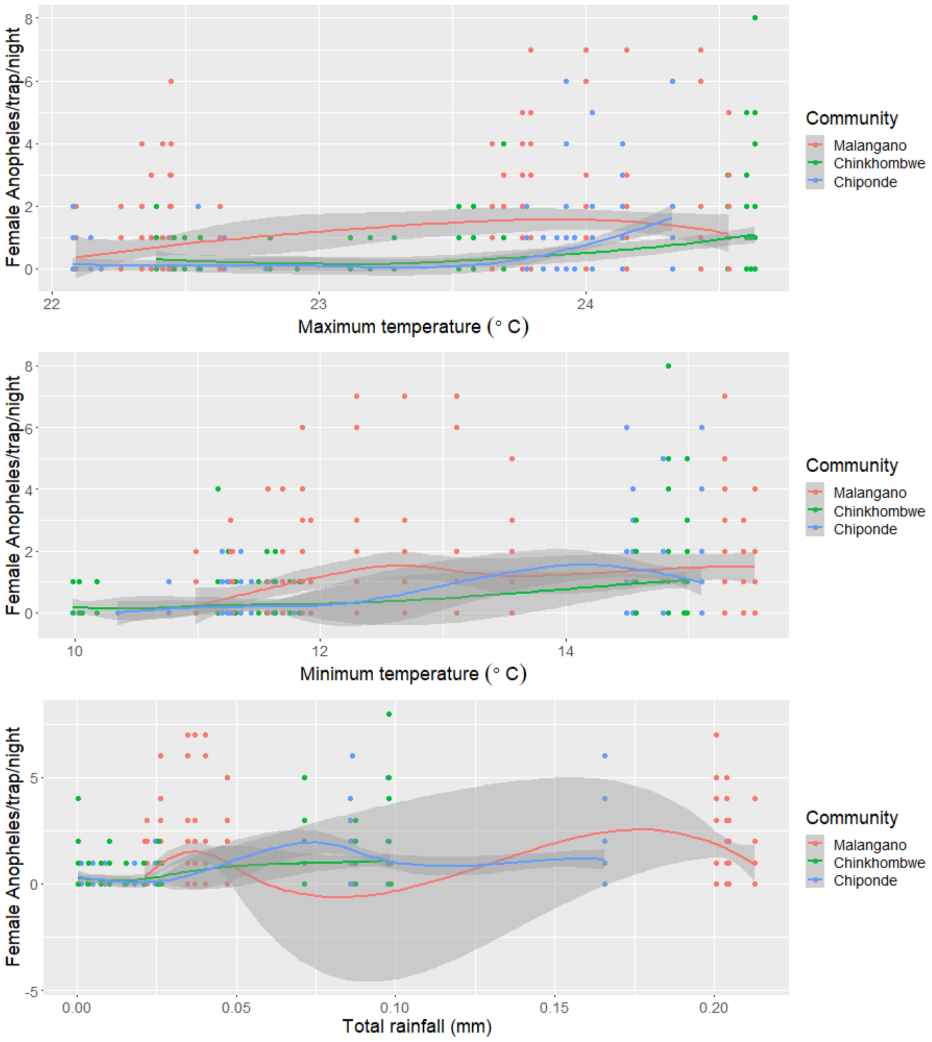

H. Model diagnostics for the best fitting model.

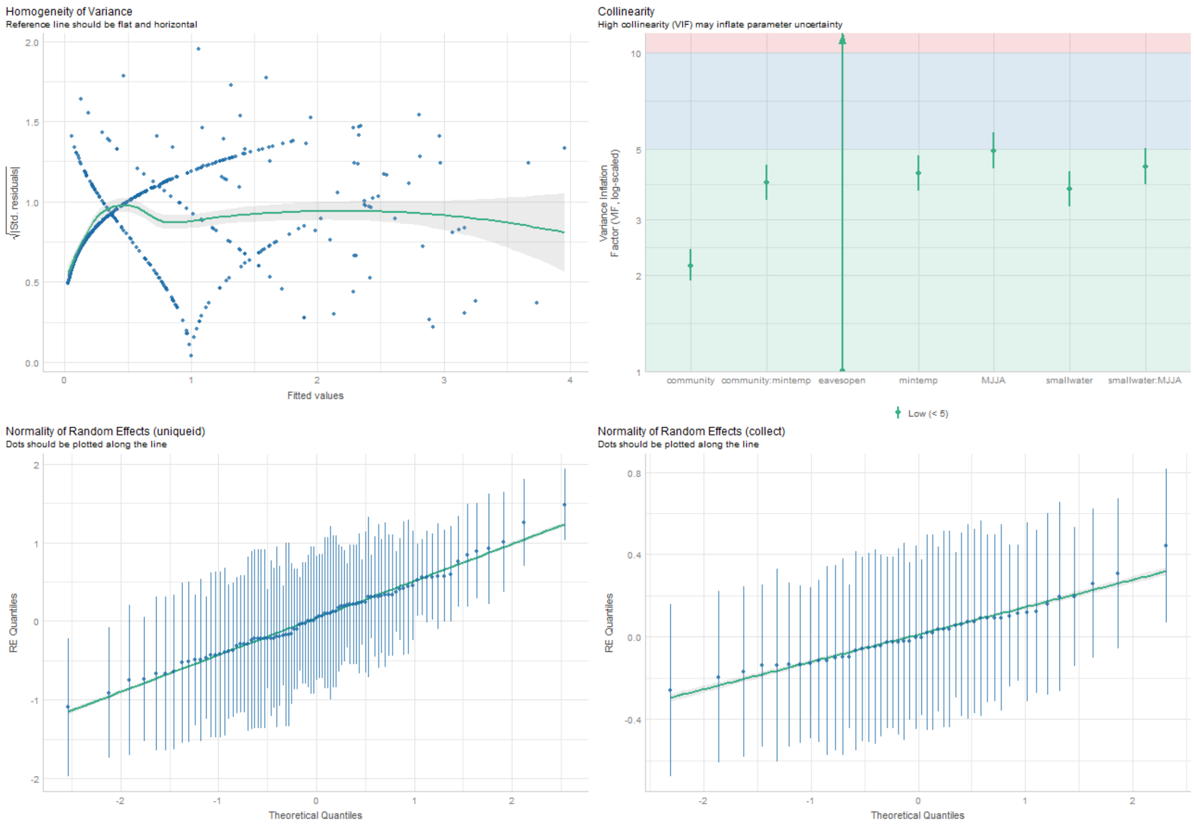

I. Boxplots of observed vs fitted female Anopheles/trap/night for the best fitting model.

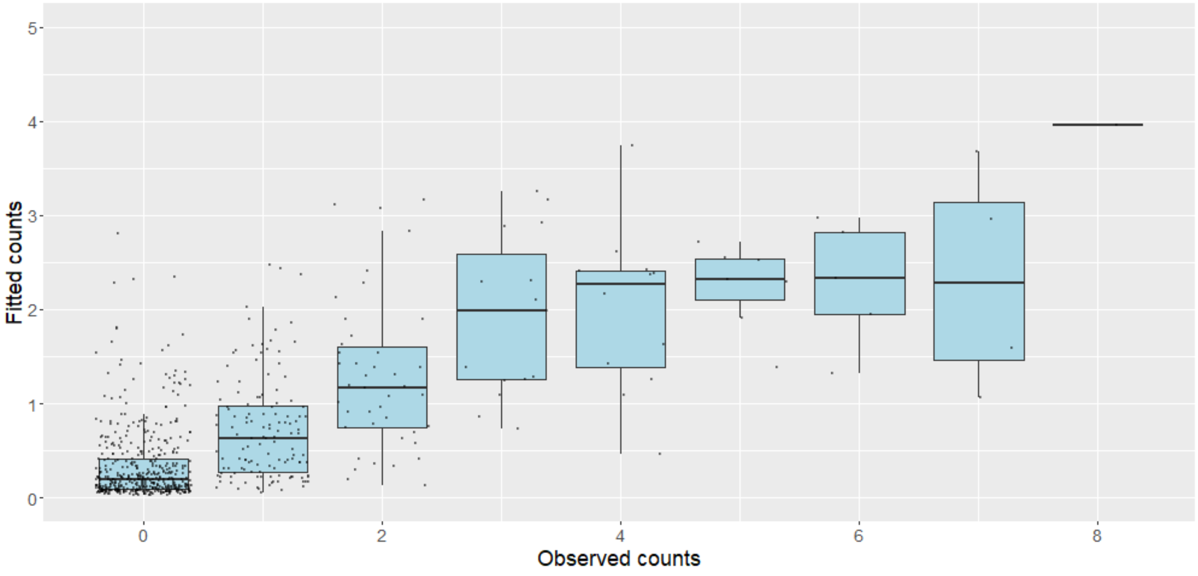
